# Ag^+^-regulated aptamers to thrombin

**DOI:** 10.1101/2025.01.09.632085

**Authors:** Rugiya Alieva, Svetlana Sokolova, Ilya Oleynikov, Roman Novikov, Timofei Zatsepin, Andrey Aralov, Elena Zavyalova

## Abstract

Nucleic acid aptamers are artificial recognition elements with high potential in biotechnology. Approaches for aptamer affinity optimization and activity regulation are required for aptamer-based nanodevices. Here thrombin aptamer affinity was increased by a single nucleotide modification with 7,8-dihydro-8-oxo-1,N^6^-ethenoadenine. Whereas a double modification makes the aptamer activity Ag^+^-dependent.

## Introduction

Nucleic acid-based aptamers are artificial oligonucleotides with specific spatial structures that are able for specific recognition of chemicals or biologicals. Aptamers have been developed to a variety of targets including inorganic cations, small organic molecules, peptides, proteins, etc.High affinity and specificity of the aptamers have been used to create pharmaceuticals and biosensors (Brown et al., 2024; He et al., 2023; Sun et al., 2023; Yin et al., 2023). Interest to this field is ongoing as oligonucleotides are much easier in handling compared to proteins. Aptamers are prone to irreversible denaturation, can be easily modified with any desirable functional group during chemical synthesis; the aptamer function can be easily tuned by a complimentary oligonucleotide or a regulatory molecule. The latest possibility is promising in development of nanodevices and artificial signal systems providing turning on / off of the aptamer activity depending on the concentration of a ligand (Zavyalova and Kopylov, 2018). For example, pH dependent i-motif-based DNA aptamer to influenza A virus has been reported recently having a high affinity at pH 6.0 and a 200-time lower affinity at pH 8.0 (Tsvetkov et al., 2024). Also, a copper (II) cation dependent DNAzyme has been developed introducing imidazole rings into the structure of natural DNAzyme providing 5-times changes in catalytic activity depending on Cu^2+^ concentration (Takezawa et al., 2024). Further expanding a basic 4-letter alphabet of natural nucleic acids with a vast variety of functional groups provides a possibility of rational design of an aptamer with specified properties.

Here we studied a potential of 7,8-dihydro-8-oxo-1,N^6^-ethenoadenine (oxo-εA) modification as a modulator of aptamer properties. The modification combines structural features of two naturally occurring lesions, 7,8-dihydro-8-oxoadenine and 1,N^6^-ethenoadenine. The modification mimics H-bonding of thymine forming a pair with adenine in DNA duplex (Figure 1A) (Aralov et al., 2022). Oxo-εA can form other types of non-canonical base pairing for example, Ag^+^-mediated oxo-εA dimer, oxo-εA–Ag^+^ _2_–oxo-εA (Figure 1B), that can stabilize antiparallel-stranded DNA duplex (Schönrath et al., 2021).

**Figure 1.**
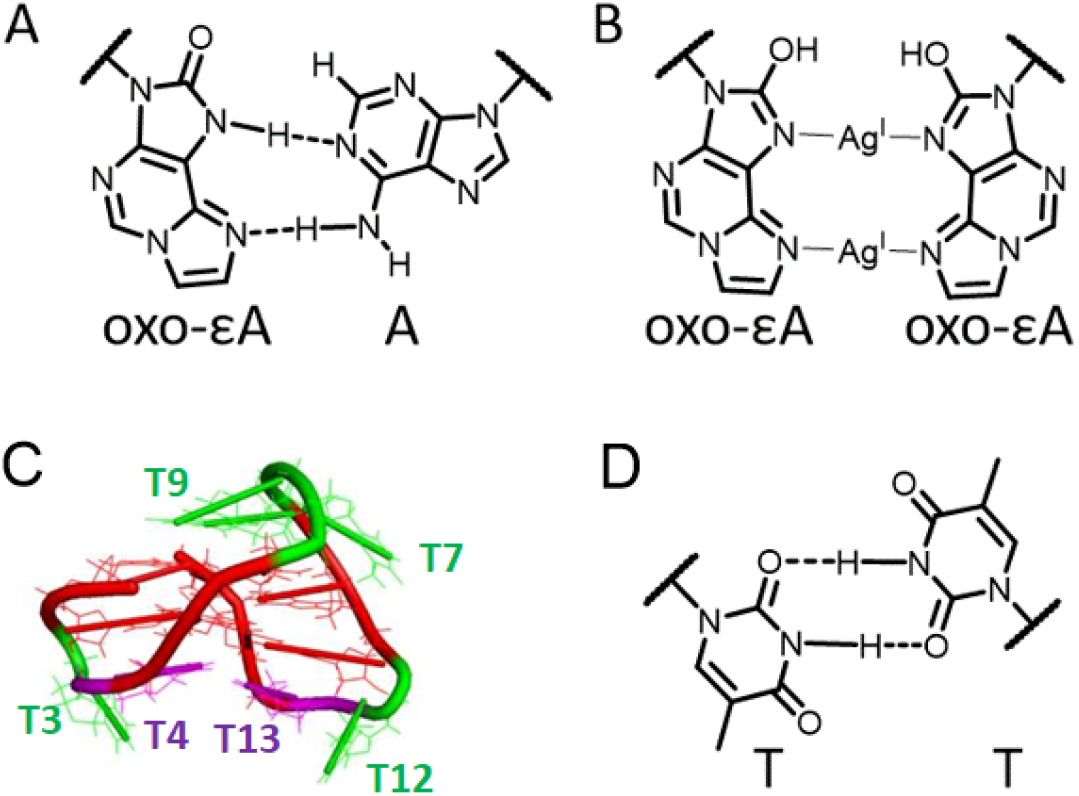
Base-pairing of 7,8-dihydro-8-oxo-1,N^6^-ethenoadenine (oxo-εA) adapted for G-quadruplex-based DNA aptamer to thrombin (HD1). Previously studied base-pairing of oxo-εA with adenine (A) and Ag^+^-mediated base-pairing between two oxo-εA modifications (B) are shown. Oxo-εA was introduced in HD1 aptamer instead of one or two thymines in the lateral loops (C). The original aptamer contains non-canonical TT-pair between T4 and T13 residues that participates in thrombin recognition (D).

G-quadruplex-based DNA aptamer to thrombin is one of the most popular objects to investigate the possibilities of rational design of aptamers. Thrombin binding aptamer (TBA, HD1) is the first identified aptamer (Figure 1C); its structure and interactions, as well as chemical modification and functional activity were reported comprehensively in dozens of papers (Riccardi et al., 2021). Rational design improved greatly the characteristics of the aptamer. The most common approaches in the rational design include chemical modifications of the heterocycles and the backbone, addition of some special structural module or combination of several aptamers in homo-or hetero-oligomeric structures. All these points are comprehensively described by Riccardi et al. (2021).

Functional activity of G-quadruplex-based aptamers is inherently metal-dependent due to the coordination of cations inside the G-quadruplex. Potassium and sodium cations promote formation of complexes with thrombin (Russo Kraus et al., 2012), whereas ammonium, barium, and strontium cations disrupts the interaction with thrombin (Zavyalova et al., 2016). In this work we studied, 8-oxoethenoA, a thymine analogue with extended aromatic structure as a modulator of thrombin aptamer affinity. The modification of T4 or T12 thymines was expected to have the most prominent increase in affinity due to stabilization of the complex with thrombin according to energy dissipating hypothesis published previously (Zhdanov et al., 2021). Additionally, we utilize the possibility of 8-oxoethenoA to form organometallic complexes with Ag^+^ to make a new type of cation-tunable aptamer. Ag^+^ cations alternate the aptamer activity due to binding to the aptamer loops involved in thrombin recognition (Figure 1D). The work describes a nice example in the outline of chemical engineering of biological recognition elements with specified properties.

## Results and Discussion

### A single modification of the T4,T13-pair improves the affinity of the aptamer to thrombin

Firstly, we introduced a single oxo-εA modification instead of one of the thymines in G-quadruplex loops (Table 1). The common approach for a new modification is to make a screening among several sites for modification introduction (Russo Kraus et al., 2012). 8 guanines of HD1 aptamer participate in formation of G-quadruplex; they were not modified to retain tertiary structure. The loops of HD1 aptamer contain 6 thymines, so 6 variants with single oxo-εA modification were studied (Table 1). The derivatives were studied with circular dichroism spectroscopy to check the integrity of G-quadruplex core and estimate melting temperatures of the structure.

**Table 1.**
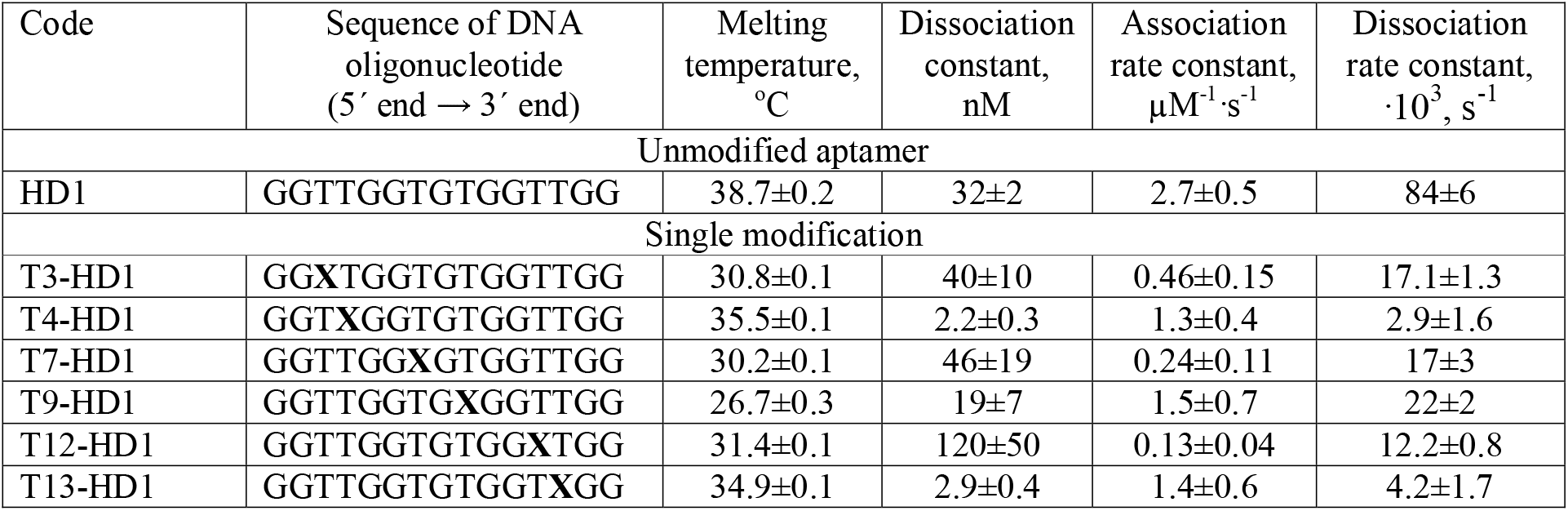
Sequences, thermal stability and affinity of thrombin aptamers modified with a single oxo-εA modification. Modification is labeled as **X**. Thermal stability was estimated with circular dichroism spectroscopy; melting temperatures were derived from the thermal dependences of circular dichroism at 295 nm. Affinity was estimated using biolayer interferometry with thrombin immobilization on the sensor.

A single modification of HD1 with oxo-εA did not alter the overall structure of G-quadruplex core (Figures S1-S3). The circular dichroism spectra are characteristic for an antiparallel G-quadruplex having two positive (247 and 295 nm) and one negative (267 nm) maxima. Thermal stability of G-quadruplex was diminished significantly by 7-12^°^C for T3-HD1, T7-HD1, T9-HD1 and T12-HD1 derivatives indicating a destabilization effect of oxo-εA on the G-quadruplex core (Table 1). Notably, T4-HD1 and T13-HD1 derivatives with oxo-εA modification within the TT-pair had much closer melting temperatures to HD1 with a difference 3-4^°^C only. The stabilization effect was accompanied with a slight distortion of the spectra. Unmodified HD1 and most of the derivatives had ratio between maxima at 295 nm and 267 nm in the range of 2.8-3.5, whereas T4-HD1 and T13-HD1 derivatives had ratios 2.1 and 2.2, correspondingly. Both maxima were larger in the case of T4-HD1 and T13-HD1 derivatives compared to HD1 aptamer, but the increase in the maximum at 267 nm was prevalent.

Next, the affinity was estimated using biolayer interferometry approach. Thrombin was immobilized onto amine-reactive sensor and allowed to interact with the aptamer solution. The sensorgrams are provided in Figures S4 and S5. The modifications differ drastically both by the signal intensities and dissociation rates of the complex. The signal intensity reflects the quantity of the assembled complex. This parameter was 10-fold higher for T4-HD1 and T13-HD1 derivatives with oxo-εA modification within the TT-pair compared to the initial HD1 aptamer and modifications of the same loops outside of the TT-pair (T3-HD1 and T12-HD1). Also, T4-HD1 and T13-HD1 aptamers had 20-30 times lower dissociation rate constants compared to the initial aptamer (Table 1).

The distinguished properties of T4-HD1 and T13-HD1 derivatives were expectable based on the previous studies. Bioinformatic analysis revealed that T3, T4, T12 and T13 residues (or their modified analogues) the hydrophobic parts of the aptamer recognizing loops are in the tight contact with the solvent in the high-affine complexes (Zhdanov et al., 2021). The extension of the solvent accessible surface of nucleotides in the complex has a stabilizing effect diminishing the dissociation rate greatly. The most pronounced effects were shown for the modifications of in the T4 and T13 positions (Zhdanov et al., 2021). The same trend can be illustrated by the data in Table 1, but the effect on the affinity is much more pronounced due to a favorable relation of dissociation and association rate constants. The equilibrium dissociation constants of T4-HD1 and T13-HD1 are among the lowest compared to other HD1 derivatives with a single modification (Riccardi et al., 2021; Zhdanov et al., 2021).

Interestingly, oxo-εA modification of T4 or T13 residues provided a slight stabilization effect on the thermal stability of G-quadruplex compared to other derivatives. The rupture of the TT-pair typically causes the total loss of affinity to thrombin and 6-8^°^C decrease in G-quadruplex melting temperatures (Smirnov and Shafer, 2000; Zavyalova and Kopylov, 2018). As oxo-εA modification of T4 or T13 residues provided an opposite effect, a pair between oxo-εA and the opposite thymine could be formed.

### Ag^+^-mediated tuning of G-quadruplex affinity to thrombin

The effects of Ag^+^ ions on G-quadruplex stability and aptamer affinity were tested. The G-quadruplex of unmodified aptamer has almost no changes in the presence of Ag^+^ ions, whereas a single modification with oxo-εA provides Ag^+^-dependent circular dichroism spectra (Figure S6). The excess of Ag^+^ ions unfolded G-quadruplexes at room temperature forming, probably, intermolecular aggregates or some destabilizing intramolecular intercations. The critical Ag^+^ concentration depends slightly on the site of modification being in the range from 3 to 8-fold excess of Ag^+^ ions (Figures S6, S7). The thermal stability of the G-quadruplexes of modified aptamers was the same with and without 2-fold excess of Ag^+^ ions that indicates two-state unfolding of G-quadruplexes without any stable intermediates (Table 2, Figures S8-S10). T4-HD1 aptamer was the single variant that was almost unfolded in the presence of 2-fold excess of Ag^+^ ions. The residual quantities of G-quadruplex provided some affinity (Table 2).

**Table 2.**
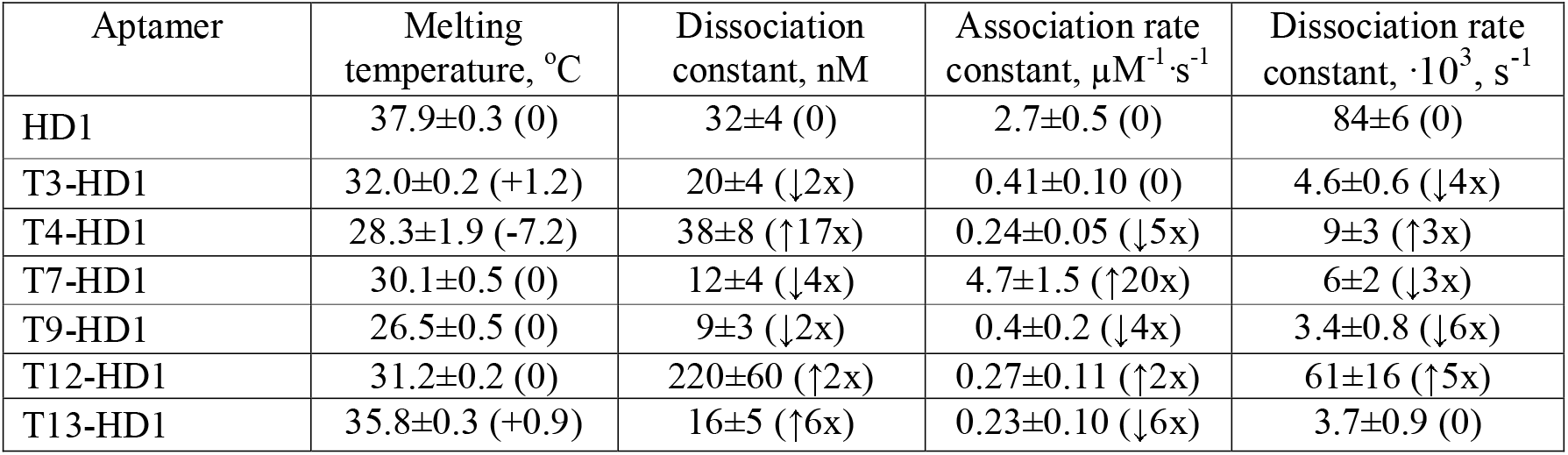
Thermal stability and affinity of thrombin aptamers modified with a single oxo-εA modification in the presence of 2-fold molar excess of Ag^+^. Thermal stability was estimated with circular dichroism spectroscopy; melting temperatures were derived from the thermal dependences of circular dichroism at 295 nm. Affinity was estimated using biolayer interferometry with thrombin immobilization on the sensor. The statistically significant changes in melting temperatures and constants compared to Ag^+^-free solution are shown in parentheses. ‘No GQ’ indicates the trace amounts of G-quadruplex in the presence of 2-fold molar excess of Ag^+^.

The dissociation constants of the aptamers with a single oxo-εA modification were determined with 5x-fold excess of Ag^+^ ions. The affinities were changed 2-17-fold, but the affinity was not abolished or greatly increased (Table 2, Figure S11). So, a single modification is not enough to make a switchable aptamer. We decided to study aptamers with dual modification.

### A dual modification of the thymine loops provides high affinity to thrombin with one exception

T3T4 and T12T13-loops were chosen for a dual modification as these loops respond for thrombin binding. 6 variants were generated (Table 3). All the aptamers retain structure of antiparallel G-quadruplex (Figures S12-S14). The melting temperatures were decreased by 1-14^°^C with one exception, T4,T13-HD1 variant that was even more stable compared to HD1 having 3^°^C increase of melting temperature (Table 3).

**Table 3.** Sequences, thermal stability and affinity of thrombin aptamers modified with a double oxo-εA modification. Modification is labeled as **X**. Thermal stability was estimated with circular dichroism spectroscopy; melting temperatures were derived from the thermal dependences of circular dichroism at 295 nm. Affinity was estimated using biolayer interferometry with thrombin immobilization on the sensor. n.d. is not determinable due to low affinity.

| Code | Sequence of DNA oligonucleotide (5' end → 3' end) | Melting temperature, °C | Dissociation constant, nM | Association rate constant, μM<sup>-1</sup>·s<sup>-1</sup> | Dissociation rate constant, ·10<sup>3</sup>, s<sup>-1</sup> |
| --- | --- | --- | --- | --- | --- |
| Unmodified aptamer |  |  |  |  |  |
| HD1 | GGTTGGTGTGGTTGG | 37.9±0.2 | 32±4 | 2.7±0.5 | 84±6 |
| Double modification |  |  |  |  |  |
| T3,T4-HD1 | GG**XX**GGTGTGGTTGG | 33.0±0.2 | 5.2±0.5 | 2.4±0.3 | 12±3 |
| T3,T12-HD1 | GG**XT**GGTGTGG**XT**GG | 24.3±1.1 | 2.2±1.1 | 0.41±0.17 | 0.75±0.09 |
| T3,T13-HD1 | GG**XT**GGTGTGGT**X**GG | 28.5±0.2 | >400 | n.d. | n.d. |
| T4,T12-HD1 | GGT**X**GGTGTGG**XT**GG | 29.8±0.3 | 1.7±0.6 | 1.4±0.3 | 2.8±1.8 |
| T4,T13-HD1 | GGT**X**GGTGTGGT**X**GG | 41.1±0.2 | >400 | n.d. | n.d. |
| T12,T13-HD1 | GGTTGGTGTGG**XX**GG | 36.5±0.3 | 6.2±1.6 | 3.0±1.3 | 17±4 |

**Table 4.** Thermal stability and affinity of thrombin aptamers modified with a dual oxo-εA modification in the presence of 2-fold molar excess of Ag^+^. Thermal stability was estimated with circular dichroism spectroscopy; melting temperatures were derived from the thermal dependences of circular dichroism at 295 nm. Affinity was estimated using biolayer interferometry with thrombin immobilization on the sensor. The statistically significant changes in melting temperatures and constants compared to Ag^+^-free solution are shown in parentheses. no GQ means the absence of maxima that are characteristic for circular dichroism spectra of antiparallel G-quadruplexes.

| Aptamer | Melting temperature, °C | Dissociation constant, nM | Association rate constant, μM<sup>-1</sup>·s<sup>-1</sup> | Dissociation rate constant, ·10<sup>3</sup>, s<sup>-1</sup> |
| --- | --- | --- | --- | --- |
| T3,T4-HD1 | 21.9±1.0 | 120±50 (↑23x) | 0.12±0.7 (↓20x) | 13±8 (0) |
| T3,T12-HD1 | no GQ | 13±3 (↑6x) | 0.22±0.02(↓2x) | 2.7±0.7 (↑4x) |
| T3,T13-HD1 | no GQ | 33±7 (↓↓) | 0.25±0.05 (↑↑) | 8±2 (↓↓) |
| T4,T12-HD1 | no GQ | 14±4 (↑8x) | 0.4±2 (↓4x) | 4±1 (0) |
| T4,T13-HD1 | 37±3 | 26±2 (↓↓) | 0.23±0.04 (↑↑) | 6.0±0.5 (↓↓) |
| T12,T13-HD1 | 38±2 | 100±50 (↑16x) | 0.11±0.06 (↓27x) | 10.6±1.3 (↓1.6x) |

T3,T13-HD1 and T4,T13-HD1 variants did not bind thrombin with high affinity, whereas all other aptamers are more affine than unmodified aptamer having dissociation constants in the range of 1.7-6.2 nM (Figure S15). The increase in affinity was acquired due to the decrease in dissociation rate constant by 5-110-fold that corresponds well with the affinity of T4-HD1 and T13-HD1 aptamers. The absence of the affinity for T3,T13-HD1 and T4,T13-HD1 variants can be explained by the formation of oxo-εA-oxo-εA pairs that are unable to recognize thrombin due to their over stabilization. T4,T13-HD1 variant seems to be more interesting for the further investigation, having unusually high thermal stability as well as the ratio between maxima at 295 nm and 267 nm of 1.6 in circular dichroism spectra that is the lowest among all studied variants.

### Ag^+^ restores the affinity of the dual modification of the T4,T13-pair

Ag^+^ ions unfolded G-quadruplexes in the stoichiometry of 1-2 (Figure S16) due to formation of oxo-εA-Ag^+^-oxo-εA pairs that are unfavorable for the G-quadruplex. Notably, T4,T13-HD1 variant is the most stable, keeping the G-quadruplex spectrum even at 3-fold excess of Ag^+^ ions. The thermal stability of T4,T13-HD1 complex with Ag^+^ ions was also the highest with melting temperature of 37^°^C that is close to the HD1 value (Table 3, Figure S17).

The affinity of T3,T4-HD1, T3,T12-HD1, T4,T12-HD1 and T12,T13-HD1 aptamers was decreased in the presence of Ag^+^ ions by 6-23-fold (Figure S18). However, the T3,T13-HD1 and T4,T13-HD1 variants with no affinity to thrombin became aptamers in the presence of Ag^+^ ions having dissociation constants nearly the same as initial HD1.

The structural reorganization was confirmed by ^1^H NMR. The chemical shifts of T4,T13-HD1 (Figure 2) were similar to HD1 shifts (Alieva et al., 2021). Addition of Ag^+^ ions altered the chemical shifts drastically (Figure 2) indicating switching of G-quadruplex conformation with preservation of its topology. The transition was completed with 2 Ag^+^ equivalents added.

**Figure 2.**
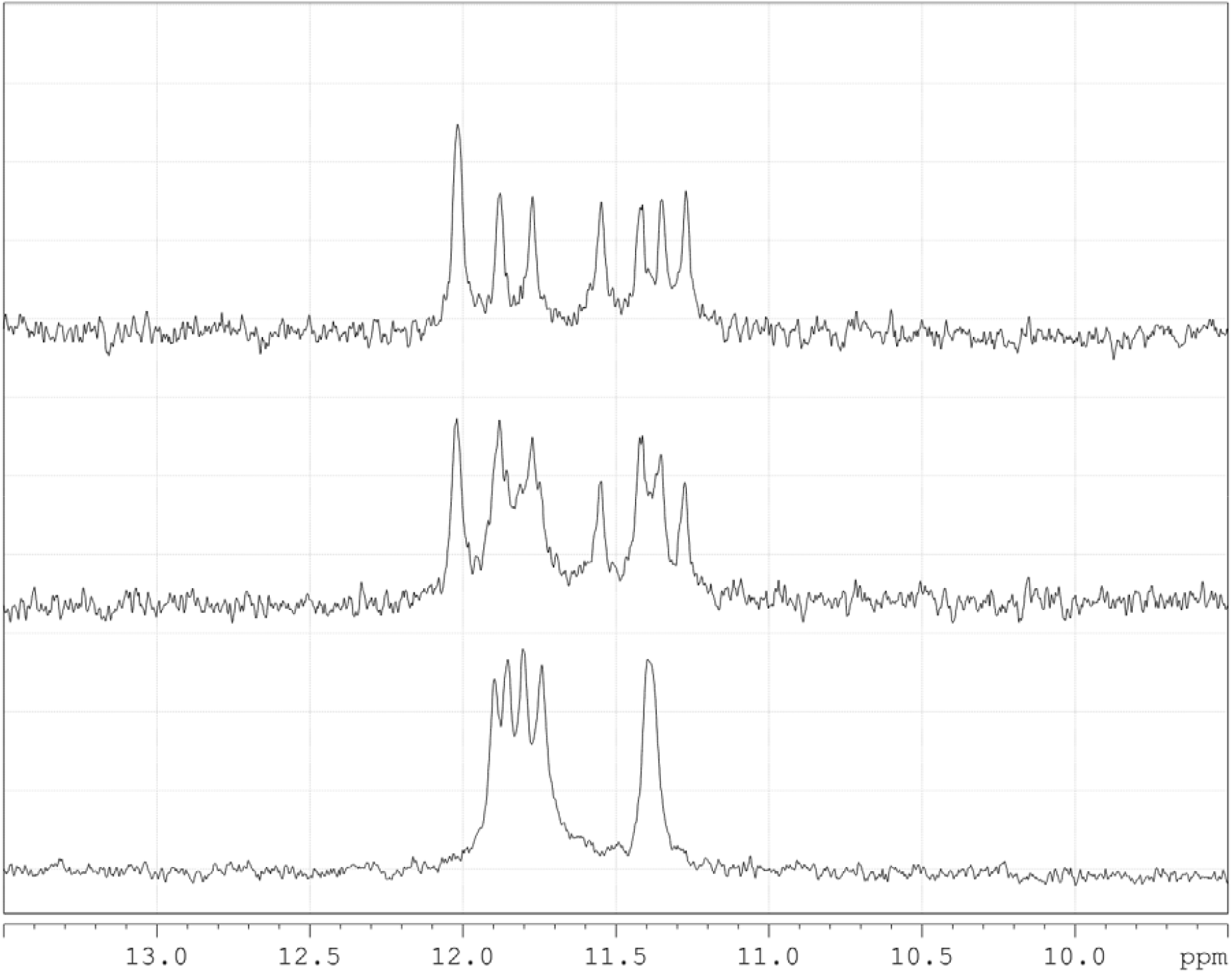
^1^H NMR spectra of T4,T13-HD1 variant with Na^+^ and K^+^ ions (the bottom spectrum), and the same solution after addition of one (the middle spectrum) and two (the upper spectrum) equivalents of Ag^+^ ions.

Summarizing, T4,T13-HD1 is a switchable aptamer with no affinity to thrombin that can be turned on by Ag^+^ ions representing a unique example of programmable recognizing elements.

## Materials and methods

### Reagents

Inorganic salts from AppliChem GmbH (Darmstadt, Germany) and Sigma-Aldrich (St. Louis, MO, USA) were used in this work. All experiments were performed in ultrapure water prepared with Milli-Q equipment (Merck KGaA, Darmstadt, Germany). HEPES buffer, pH 7.5, from AppliChem (Darmstadt, Germany), carboxymethylethanolamine (CME) from Merck (Schuchardt, Germany), and sodium salt of N-hydroxysulfosuccinimide (s-NHS) from Chem-Impex Int’l (Wood Dale, IL, USA) were used. Recombinant thrombin was purchased from Haemtech Inc. (USA).

### Oligonucleotides

Oligonucleotides were synthesized using commercially available reagents by a solid-phase phosphoramidite method, followed by purification with high performance liquid chromatography. The sequences and modification sites are shown in Tables 1 and 3. G-quadruplexes were assembled in 2 μM aptamer concentration in the buffer with the sequential heating at 95^°^C for 5 min and cooling at room temperature.

### Circular dichroism and UV spectroscopy

2 μM aptamer solutions in the buffer (10 mM Tris-HCl, pH 7.3, 140 mM NaNO_3_, 10 mM KNO_3_) were mixed with different amounts of Ag^+^ ions. The final concentrations of Ag^+^ ions were in the range of 0-64 µM. The samples were studied after short incubation at room temperature (5 minutes). The samples were placed in quartz cuvettes with a 1 cm path. Circular dichroism and UV spectra were acquired using the Chirascan spectrometer (Applied Photophysics, Leatherhead, Great Britain) and MOS-500 spectrometer (BioLogic, Seyssinet-Pariset, France) equipped with a thermoelectric temperature regulator. The spectra were acquired in the wavelength range of 240–360 nm. The spectrum of the buffer was subtracted as a baseline. The vµsamples were heated with a mean ramp of 1.0^°^C/min. The melting experiments were conducted in the range of 10–90^°^C.

### Affinity assay

The affinity of the aptamers to thrombin was estimated using biolayer interferometry (Blitz, ForteBIO, Menlo Park, CA, USA). The experiments were conducted at 20^°^C. Samples were placed in black 0.5 mL tubes (Sigma-Aldrich, St. Louis, MO, USA) in a 300 μL volume.

Biosensors intended for the amine coupling reaction (AR2G biosensors, ForteBio, Menlo Park, CA, USA) were hydrated for 10 minutes in water. The sensors were activated for 5 min in the solution of 200 mM CME and 100 mM s-NHS. Thrombin was diluted to 2.5 mg/mL in 40 mM phosphate buffer pH 6.4. The sensor was washed and blocked with 100 mM HEPES-HCl buffer with pH 7.5 for 3 min. After the signal stabilization in the buffer (10 mM Tris-HCl, 140 mM NaCl, 10 mM KCl). Aptamers were assembled in the same buffer. The association step was conducted with aptamer solution in the concentration 2000, 1000, 500, or 250 nM in the buffer. Then the dissociation step was conducted in the buffer. Experiments with Ag^+^ ions were conducted in the same buffer with addition of 5 µM of AgNO_3_. 2 μM solution of the assembled aptamer in the buffer was incubated with 5 µM of AgNO_3_ prior the experiment.

### NMR experiment

Solutions of 200 µM of T4,T13-HD1 variant was prepared in 140 mM NaNO_3_, 10 mM KNO_3_ solution, annealed for 5 minutes at 95^°^C and cooled at room or at +5^°^C temperature. The samples were diluted by D_2_O (final D_2_O content was 10%). ^1^H NMR spectra were recorded using Bruker AVANCE III HD 300, Bruker AVANCE III HD 400, and Bruker AVANCE III 600 spectrometers (Bruker, USA). ^1^H chemical shifts were referenced relative to external sodium 2,2-dimethyl-2-silapentane-5-sulfonate (DSS). ^1^D proton (^1^H) spectra of samples in H_2_O+D_2_O (10%) were recorded at +20^°^C using pulsed-field gradient ‘WATERGATE W5’ pulse sequence (zggpw5 from Bruker library) for complete H_2_O-signal clear suppression. A relaxation delay of 1.5–2.5 s and an acquisition time of 1.35–2.40 sec were used for all experiments. For a precise baseline correction of imino-protons region of spectra to suppress ‘mathematically’ a wide oligomeric hump (1H ‘oligomers filtered’ NMR spectra), ‘multipoint baseline correction Whittaker’ algorithm was applied using ‘Bruker TopSpin 3.6’ software (Bruker, Billerica, MA, USA) and ‘MestreNova 6.0’ software (Mestrelab, Spain), in addition to initial collected data with applying of the T2-relaxation-filter to partial direct suppress of oligomers during acquisition.

