## Supplementary materials for "Ag^+^-regulated aptamers to thrombin"

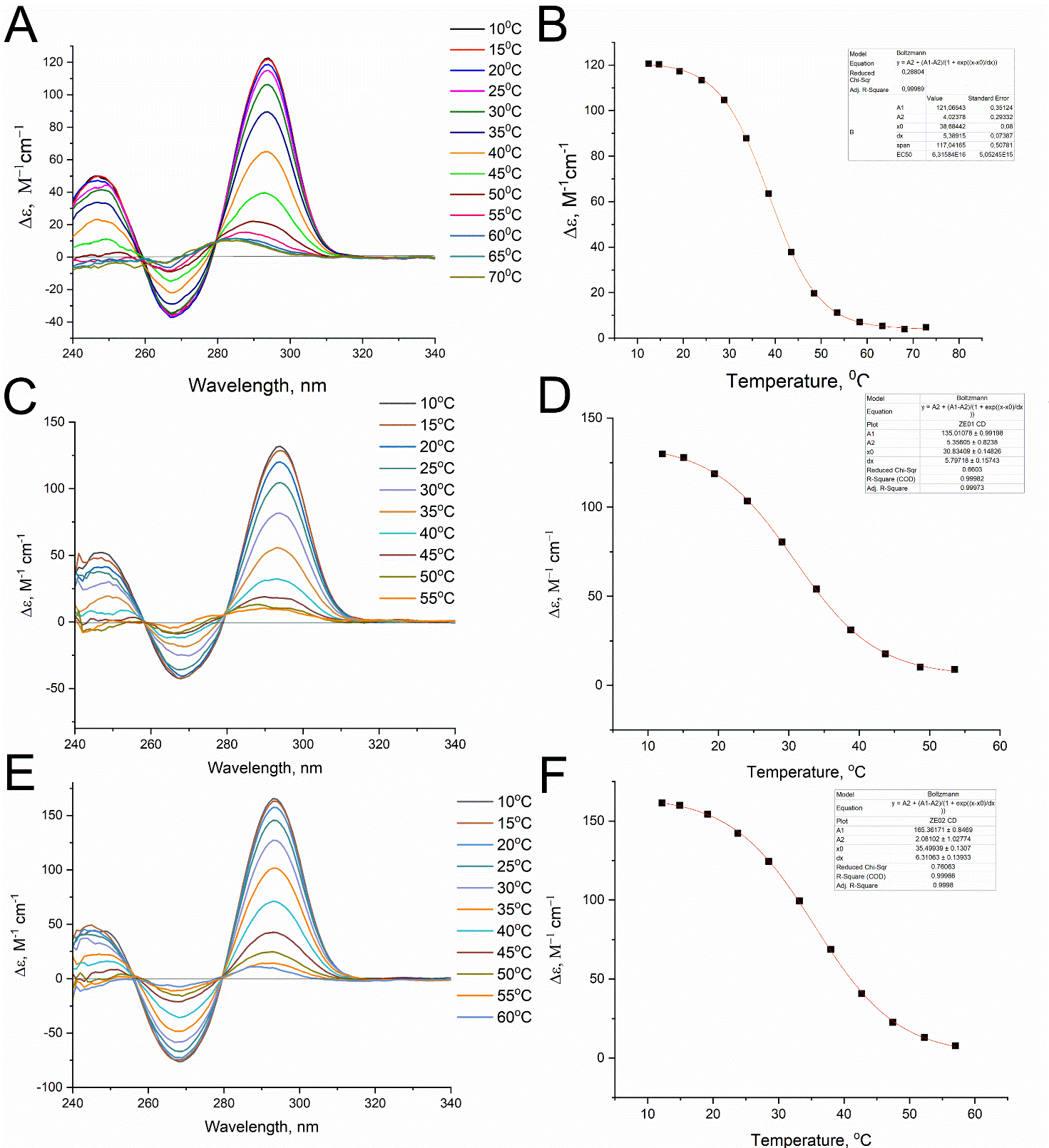

Figure S1. Thermal stability of G-quadruplexes of HD1 (A), T3-HD1 (C), and T4-HD1 (E) aptamers studied using circular dichroism spectroscopy. The melting profiles of HD1 (B), T3-HD1 (D), and T4-HD1 (F) aptamers at a wavelength of 295 nm are provided.

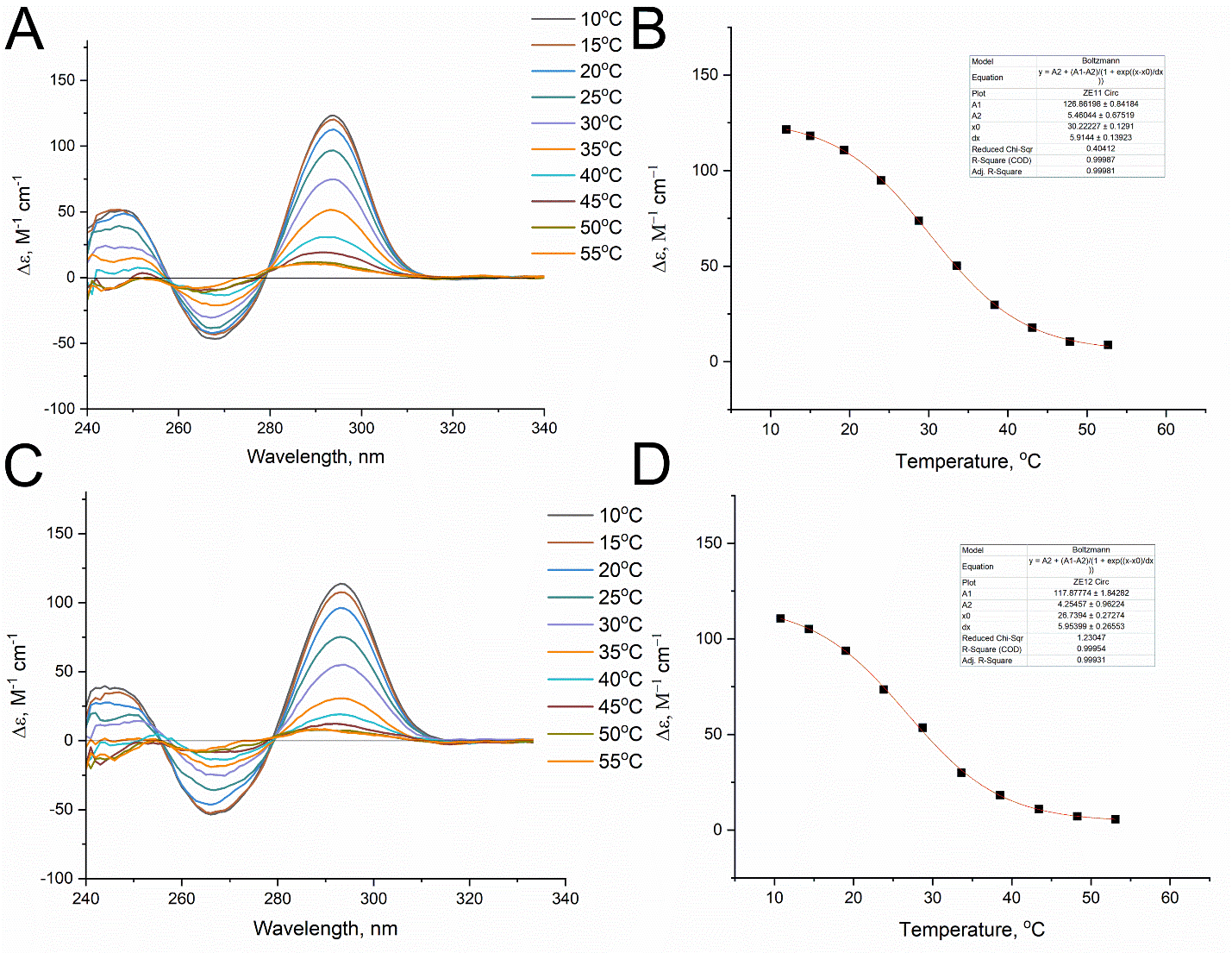

Figure S2. Thermal stability of G-quadruplexes of T7-HD1 (A) and T9-HD1 (C) aptamers studied using circular dichroism spectroscopy. The melting profiles of T7-HD1 (B) and T9-HD1 (D) aptamers at a wavelength of 295 nm are provided.

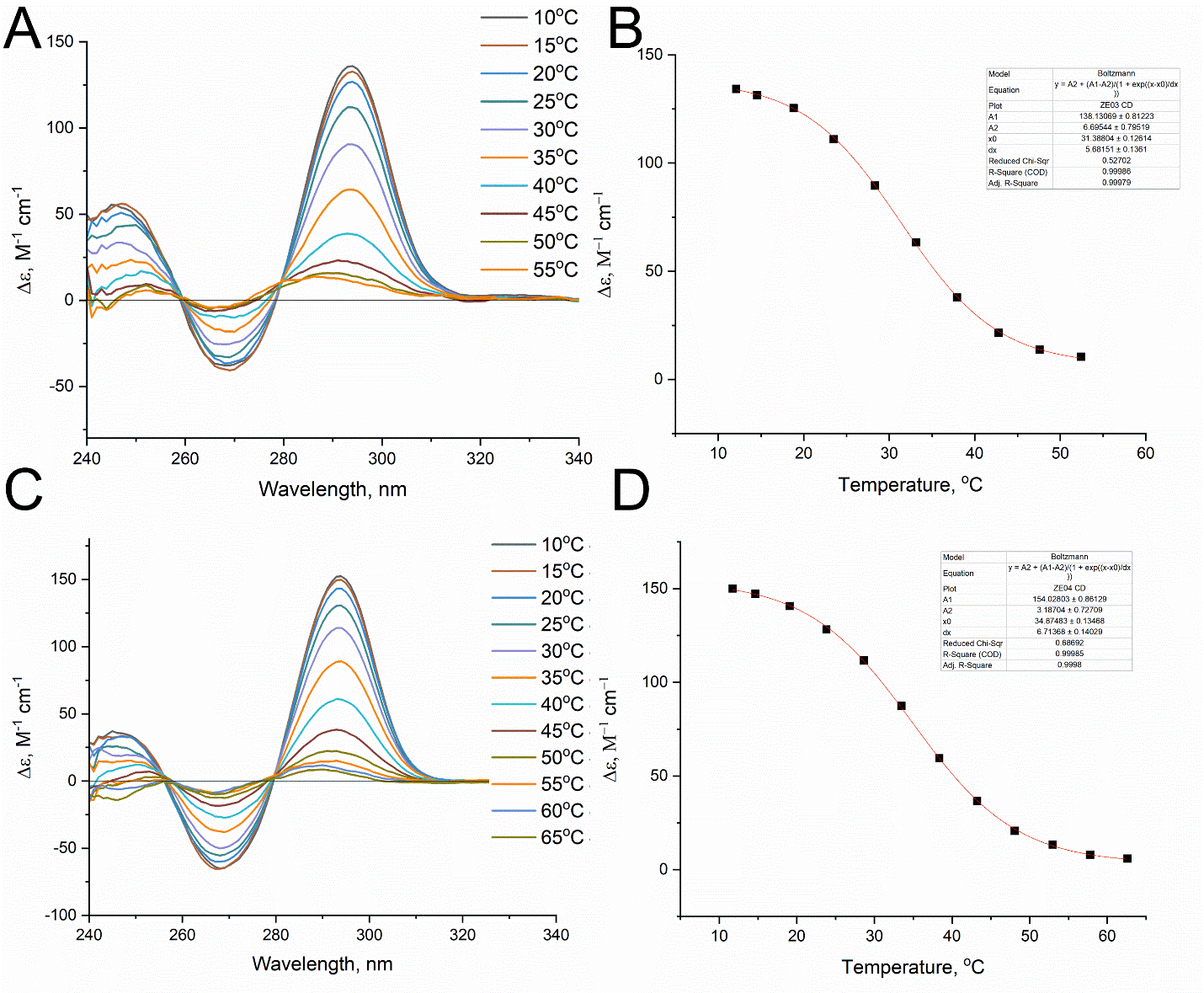

Figure S3. Thermal stability of G-quadruplexes of T12-HD1 (A) and T13-HD1 (C) aptamers studied using circular dichroism spectroscopy. The melting profiles of T12-HD1 (B) and T13-HD1 (D) aptamers at a wavelength of 295 nm are provided.

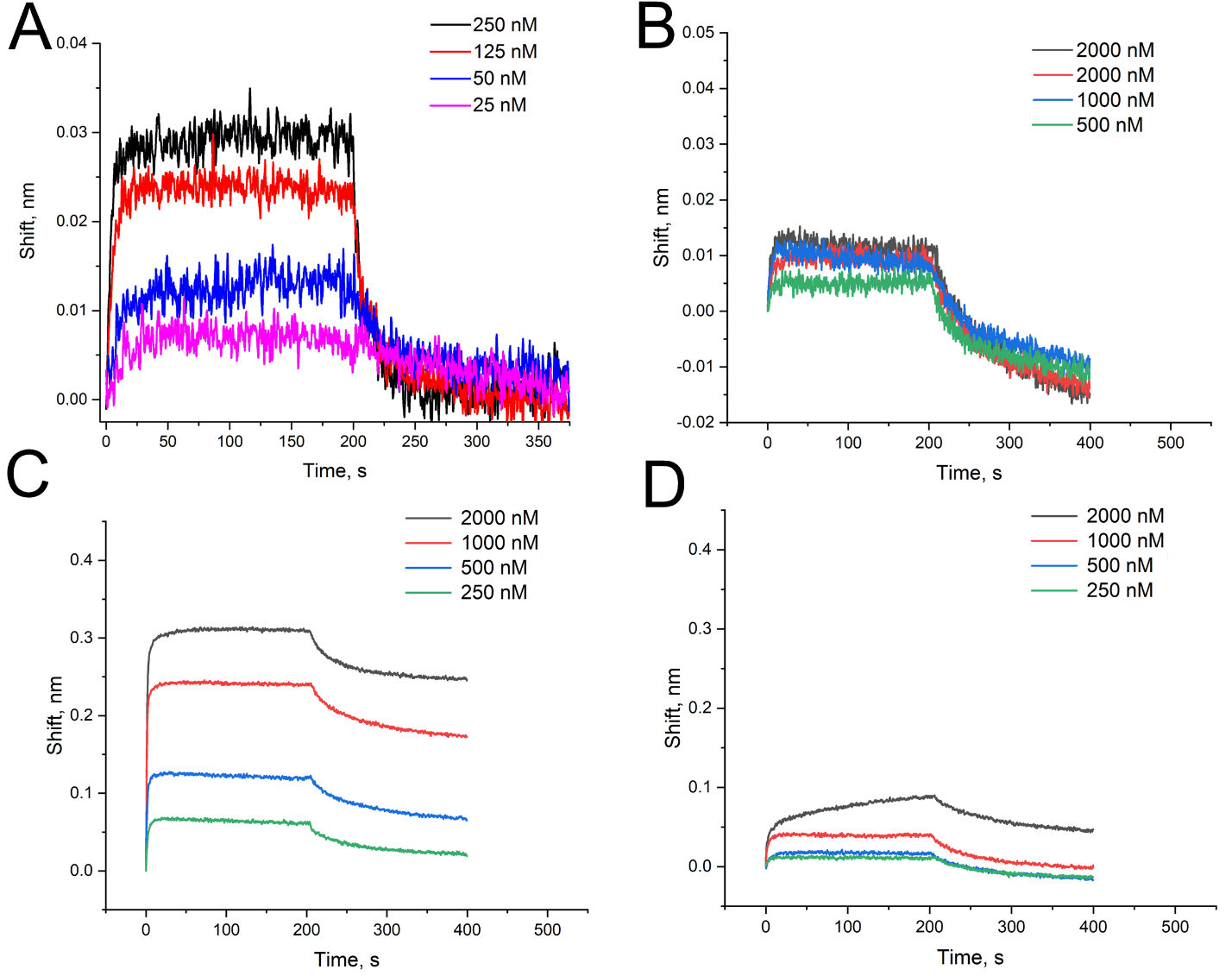

Figure S4. Biolayer interferometry sensorgrams for HD1 (A), T3-HD1 (B), T4-HD1 (C) and T7-HD1 (D) aptamers binding to immobilized thrombin. The aptamer-thrombin association stage was performed from 0 to 200 s; the aptamer-thrombin dissociation stage was performed from 200 to 400 s. The data for HD1 aptamer were adopted from (Zhdanov et al., 2021).

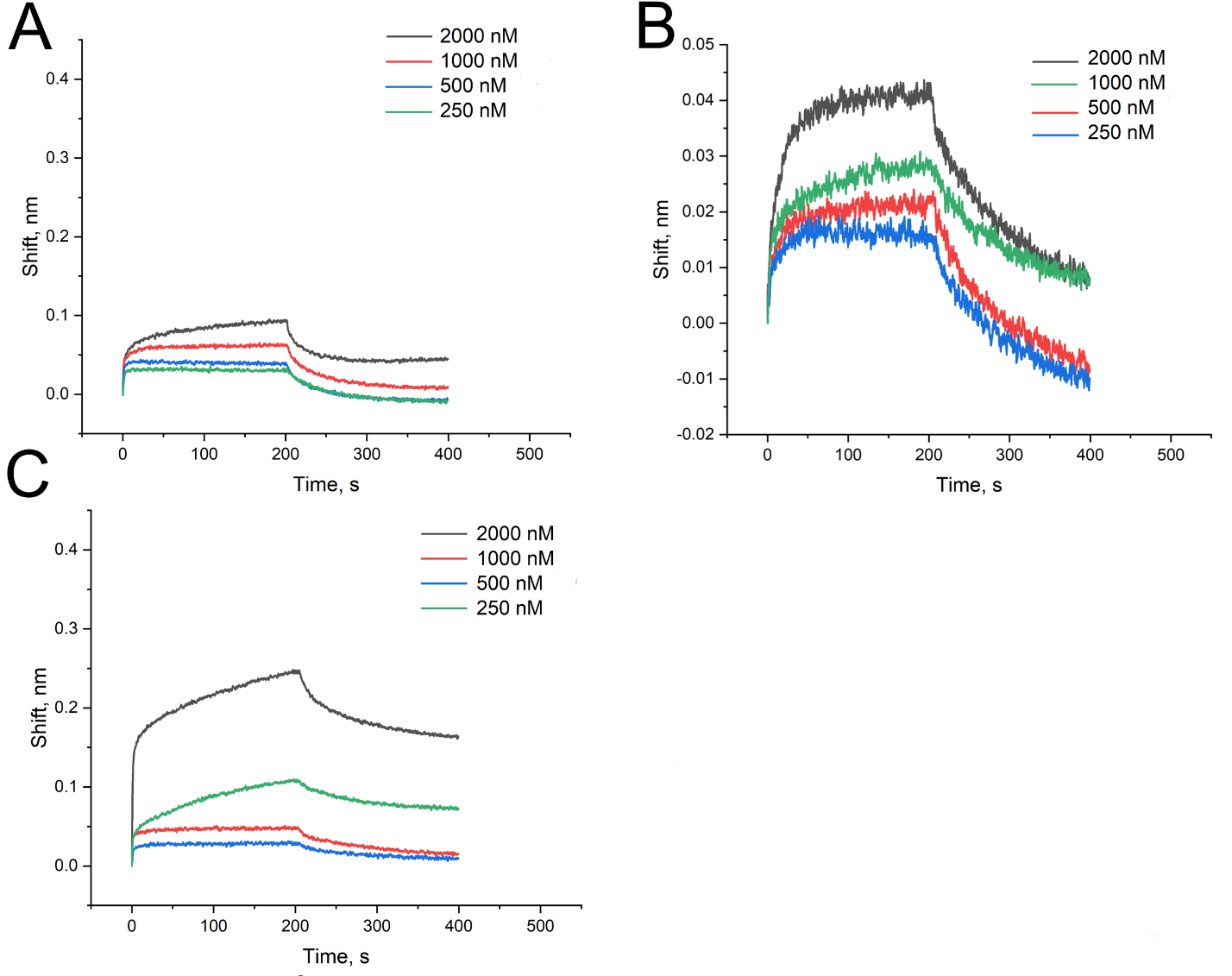

Figure S5. Biolayer interferometry sensorgrams for T9-HD1 (A), T12-HD1 (B) and T13-HD1 (C) aptamers binding to immobilized thrombin. The aptamer-thrombin association stage was performed from 0 to 200 s; the aptamer-thrombin dissociation stage was performed from 200 to 400 s.

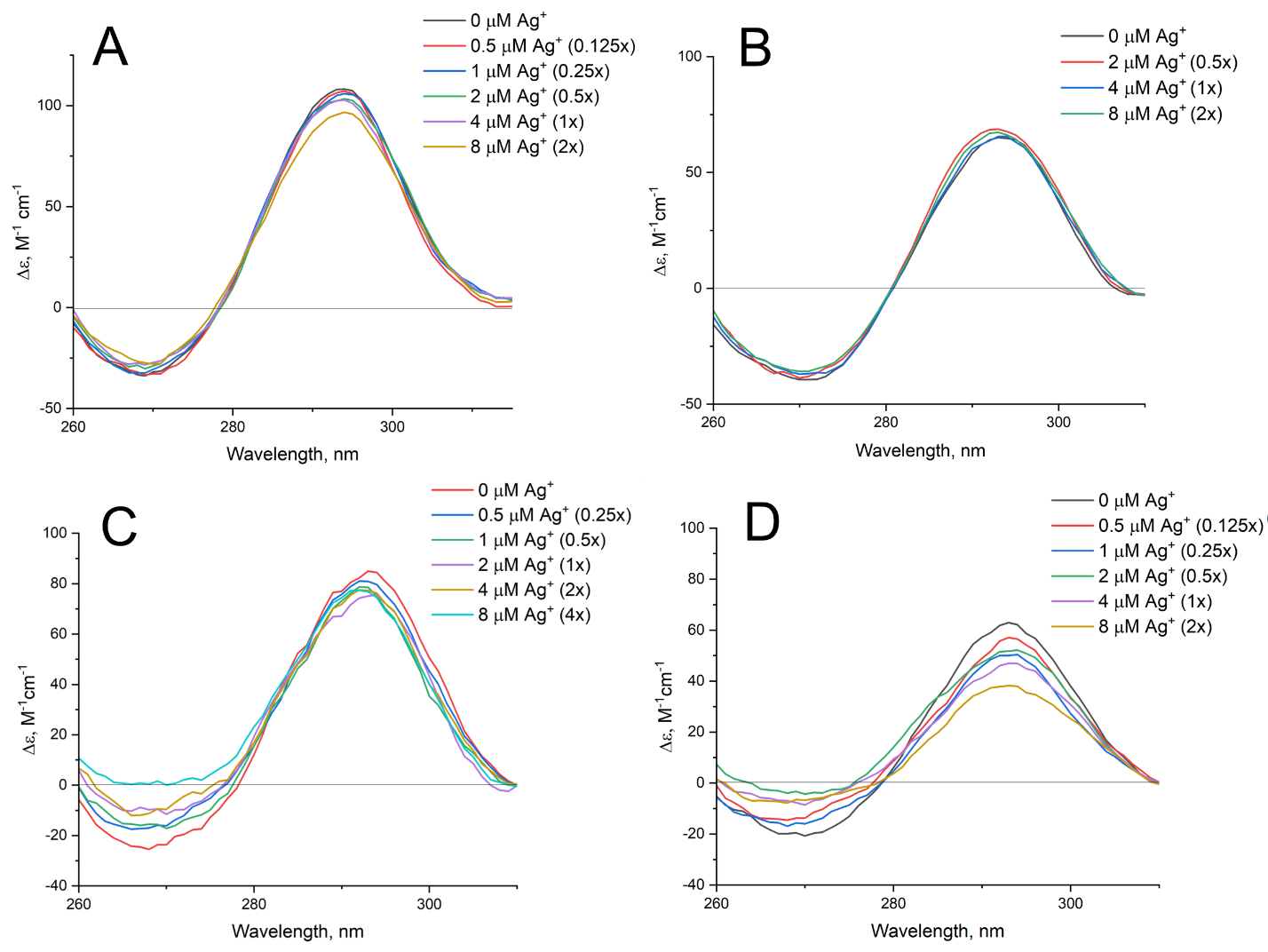

Figure S6. Circular dichroism spectra of aptamers with a single oxo-εA modification in the presence of different amounts of Ag^+^ ions. HD1 (A), T3-HD1 (B), T4-HD1 (C) and T7-HD1 (D) aptamers were assembled in the buffer, mixed with Ag^+^ for 5 minutes and placed in the spectrometer.

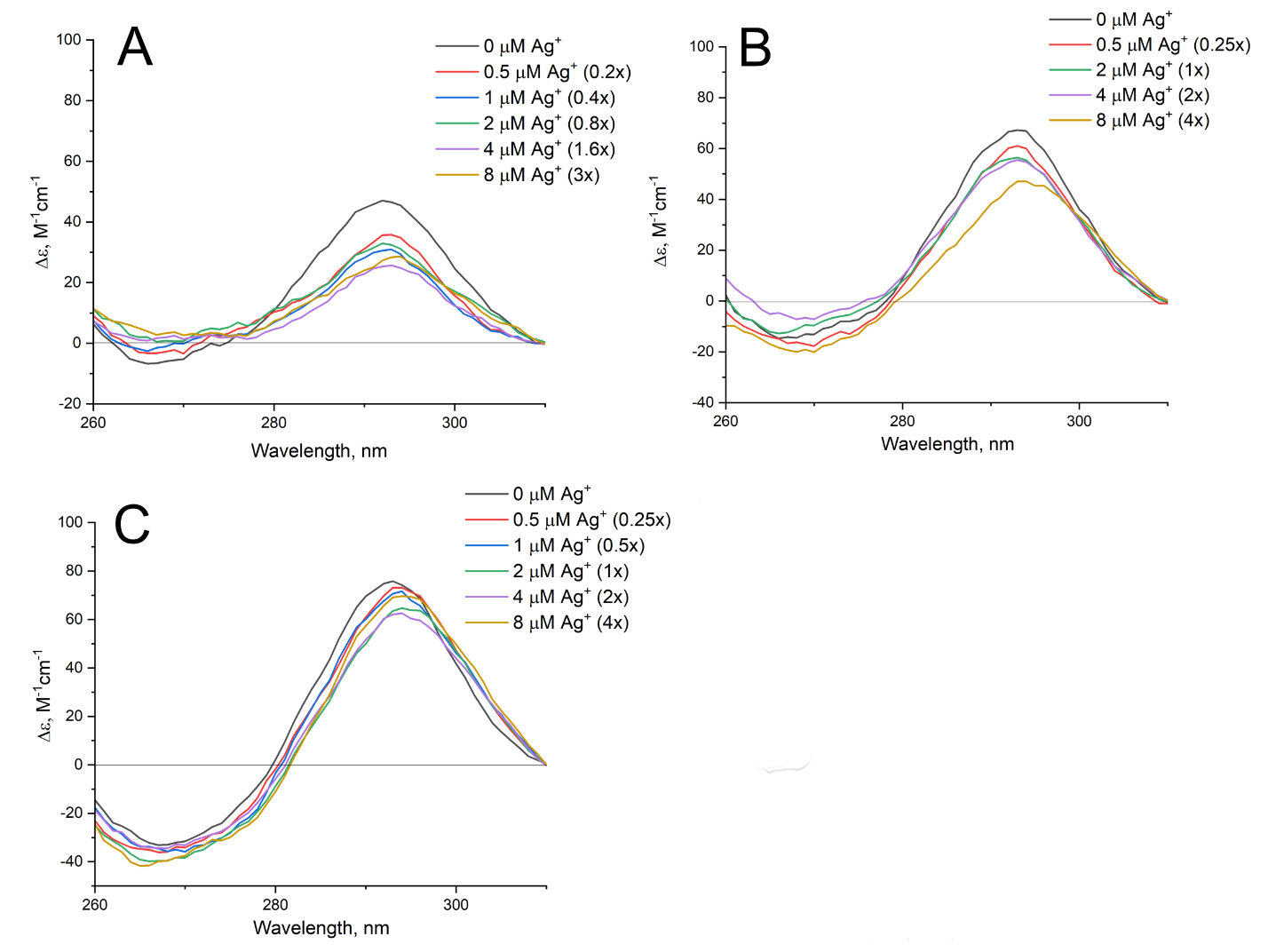

Figure S7. Circular dichroism spectra of aptamers with a single oxo-εA modification in the presence of different amounts of Ag^+^ ions. T9-HD1 (A), T12-HD1 (B) and T13-HD1 (C) aptamers were assembled in the buffer, mixed with Ag^+^ for 5 minutes and placed in the spectrometer.

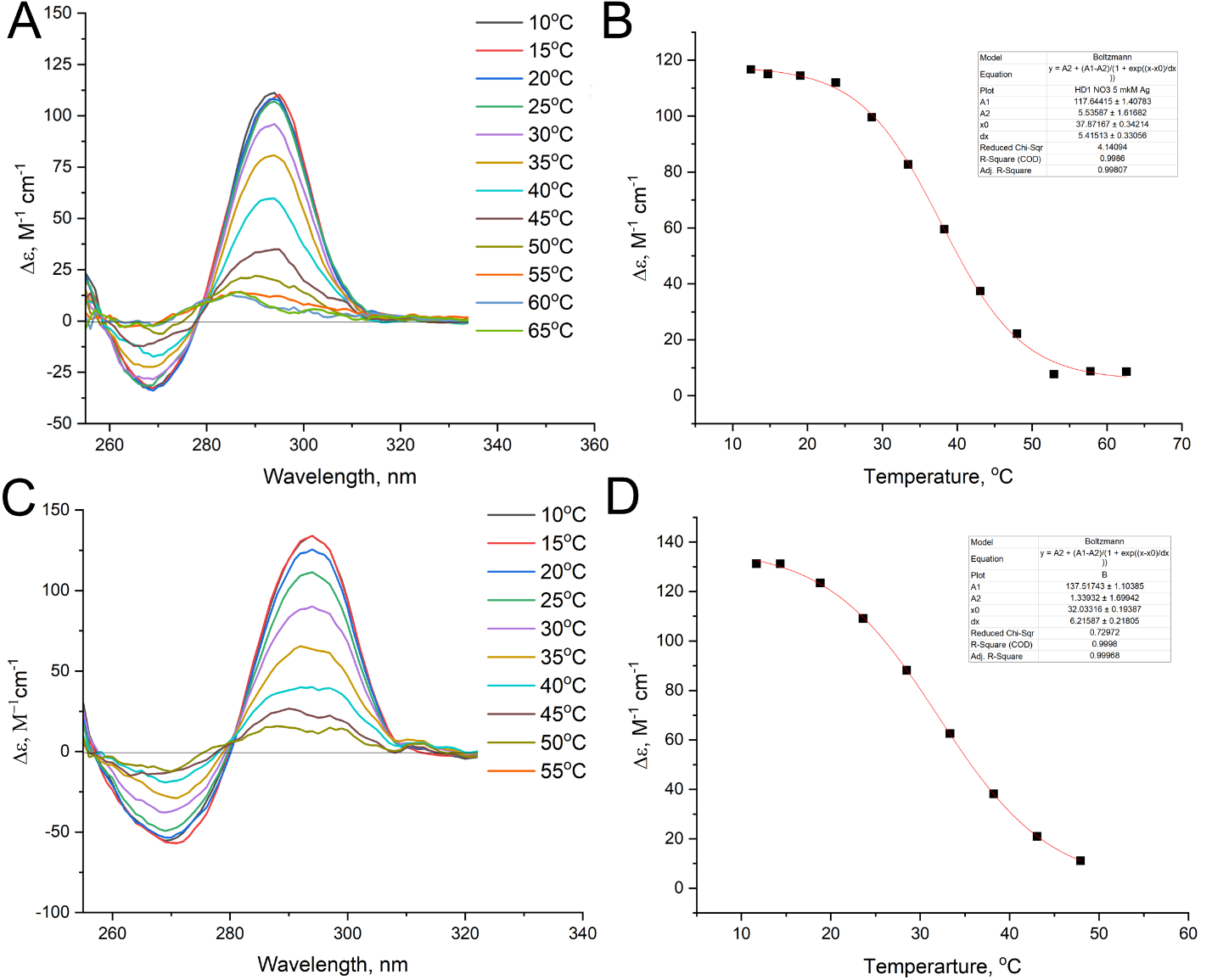

Figure S8. Thermal stability of G-quadruplexes of HD1 (A) and T3-HD1 (C) aptamers with 2-fold excess of Ag^+^ studied using circular dichroism spectroscopy. The melting profiles of HD1 (B) and T3-HD1 (D) aptamers at a wavelength of 295 nm are provided.

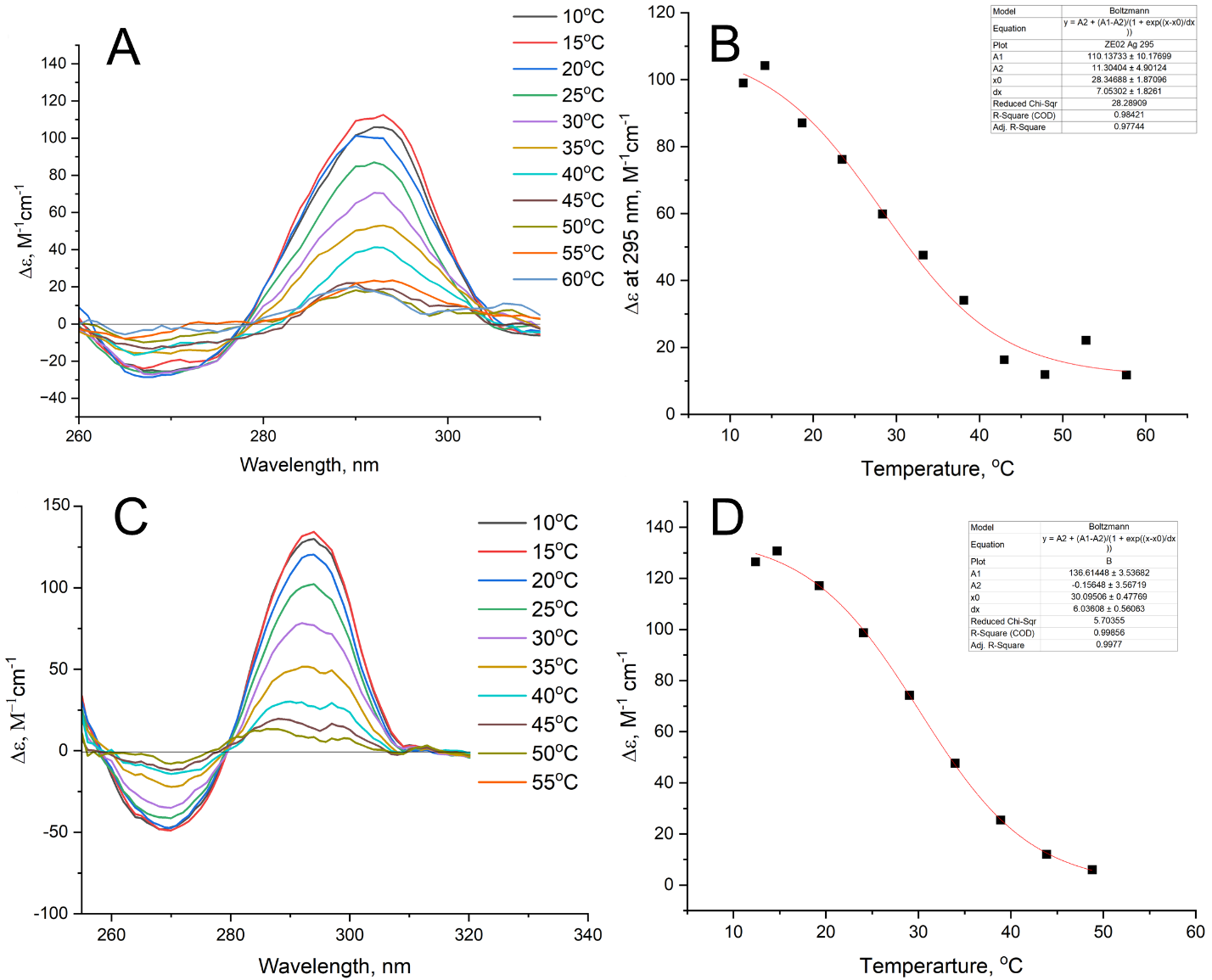

Figure S9. Thermal stability of G-quadruplexes of T4-HD1 (A) and T7-HD1 (C) aptamers with 2-fold excess of Ag^+^ studied using circular dichroism spectroscopy. The melting profiles of T4-HD1 (B) and T7-HD1 (D) aptamers at a wavelength of 295 nm are provided.

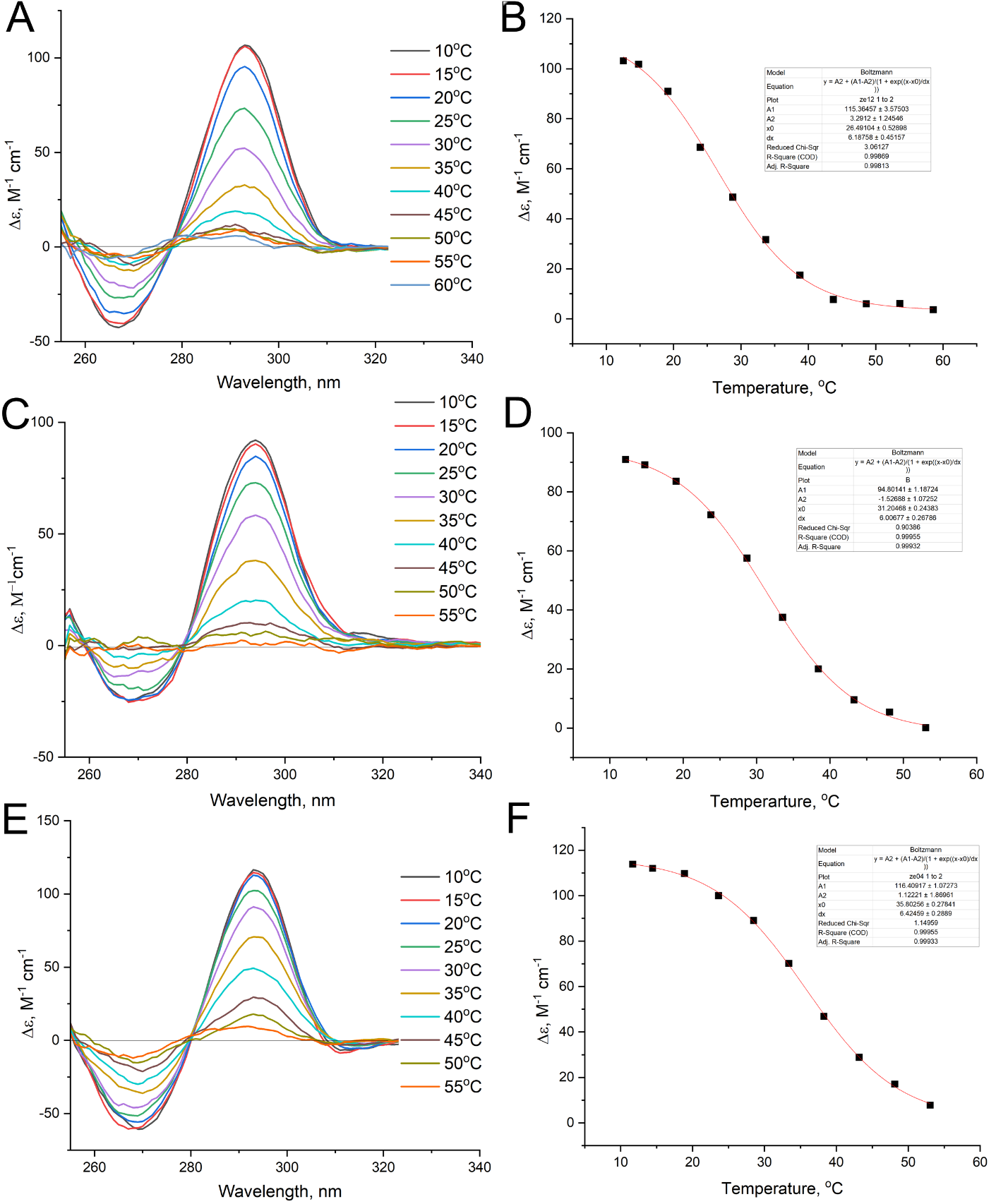

Figure S10. Thermal stability of G-quadruplexes of T9-HD1 (A), T12-HD1 (C) and T13-HD1 (E) aptamers with 2-fold excess of Ag^+^ studied using circular dichroism spectroscopy. The melting profiles of T9-HD1 (B), T12-HD1 (D) and T13-HD1 (F) aptamers at a wavelength of 295 nm are provided.

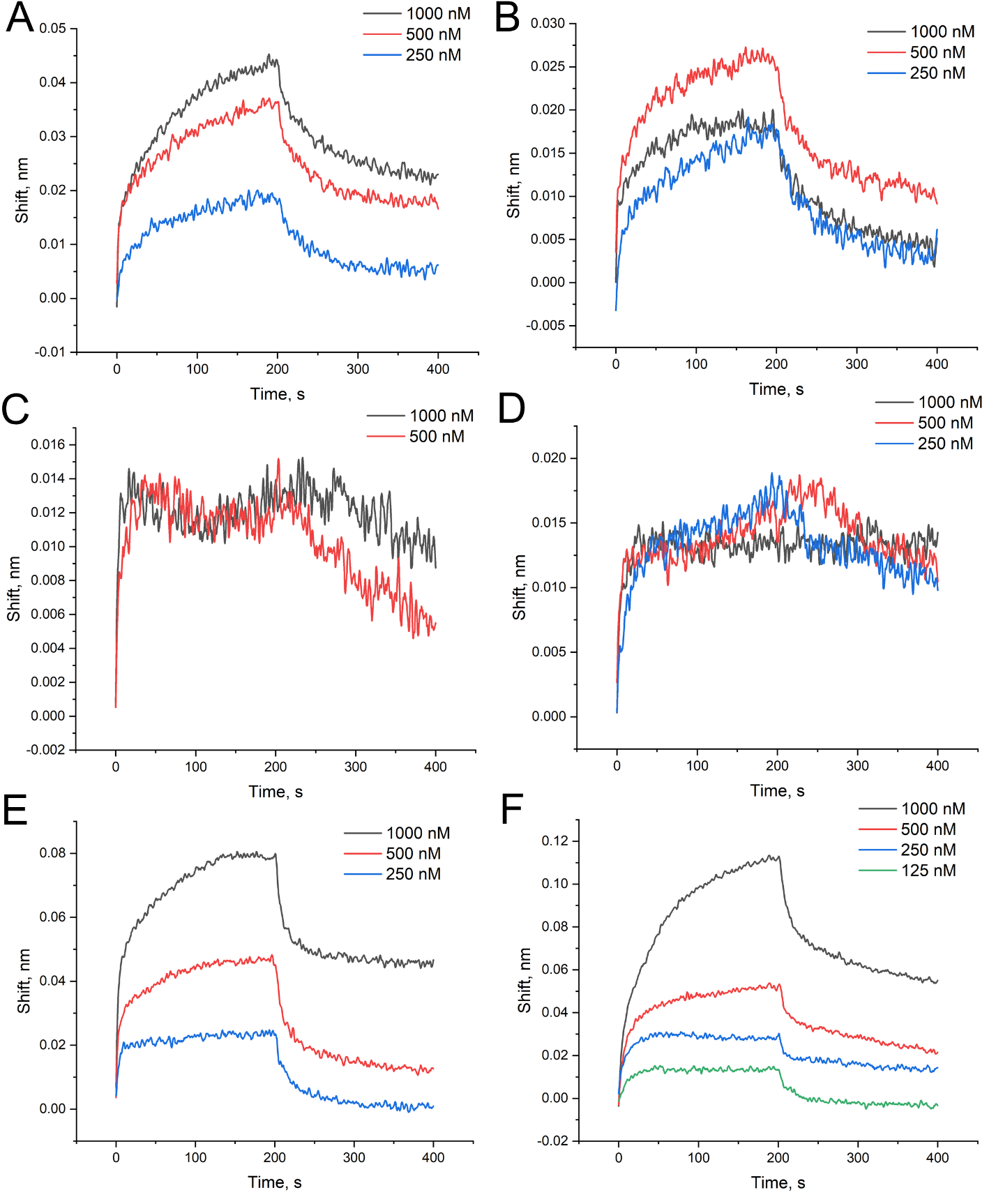

Figure S11. Biolayer interferometry sensorgrams for T3 -HD1 (A), T4-HD1 (B), T7-HD1 (C), T9-HD1 (D), T12-HD1 (E), T13-HD1 (F) aptamers with 5-fold excess of Ag^+^ that bind immobilized thrombin. The aptamer-thrombin association stage was performed from 0 to 200 s; the aptamer-thrombin dissociation stage was performed from 200 to 400 s.

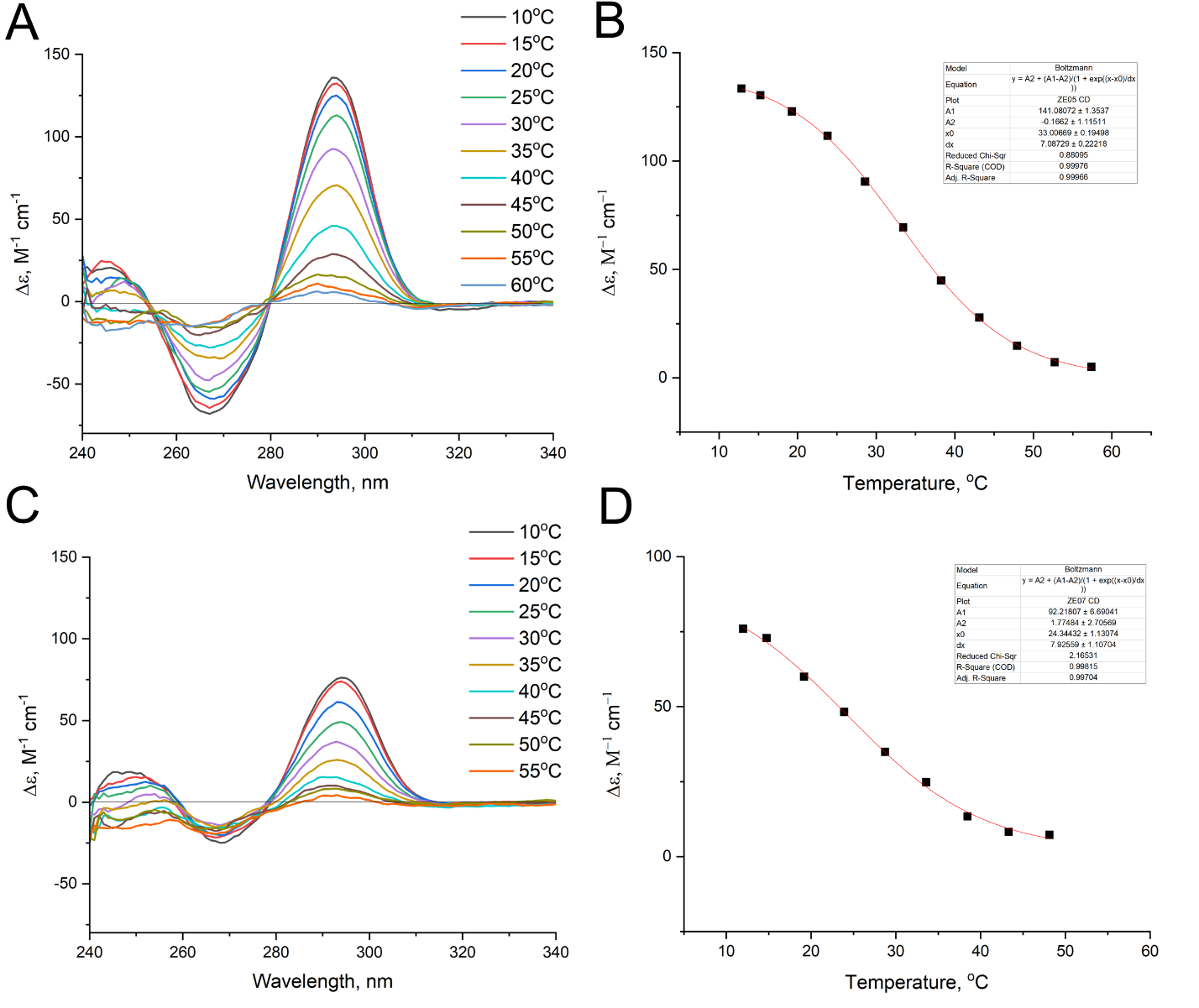

Figure S12. Thermal stability of G-quadruplexes of T3,T4-HD1 (A) and T3,T12-HD1 (C) aptamers studied using circular dichroism spectroscopy. The melting profiles of T3,T4-HD1 (B) and T3,T12-HD1 (D) aptamers at a wavelength of 295 nm are provided.

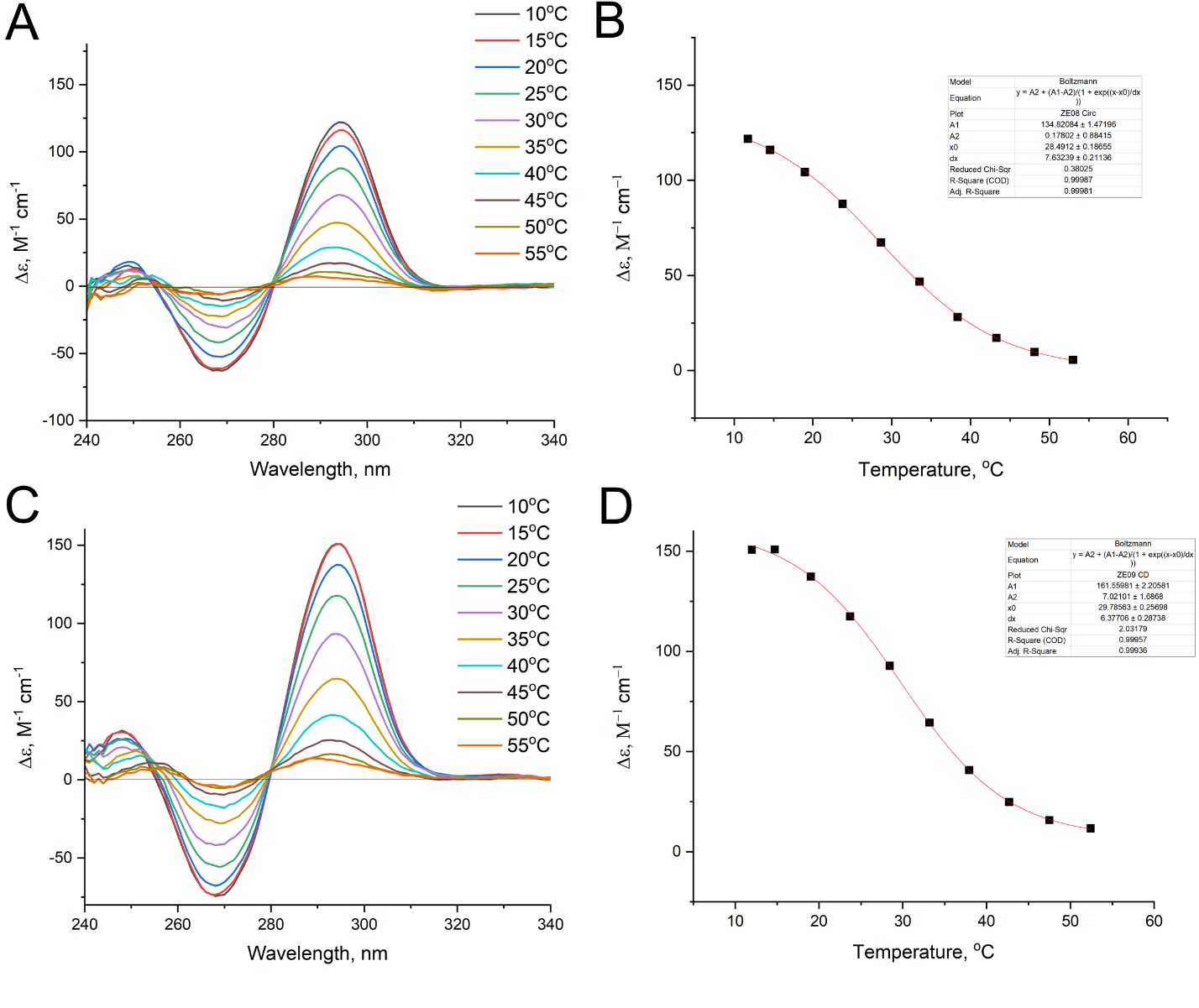

Figure S13. Thermal stability of G-quadruplexes of T3,T13-HD1 (A) and T4,T12-HD1 (C) aptamers studied using circular dichroism spectroscopy. The melting profiles of T3,T13-HD1 (B) and T4,T12-HD1 (D) aptamers at a wavelength of 295 nm are provided.

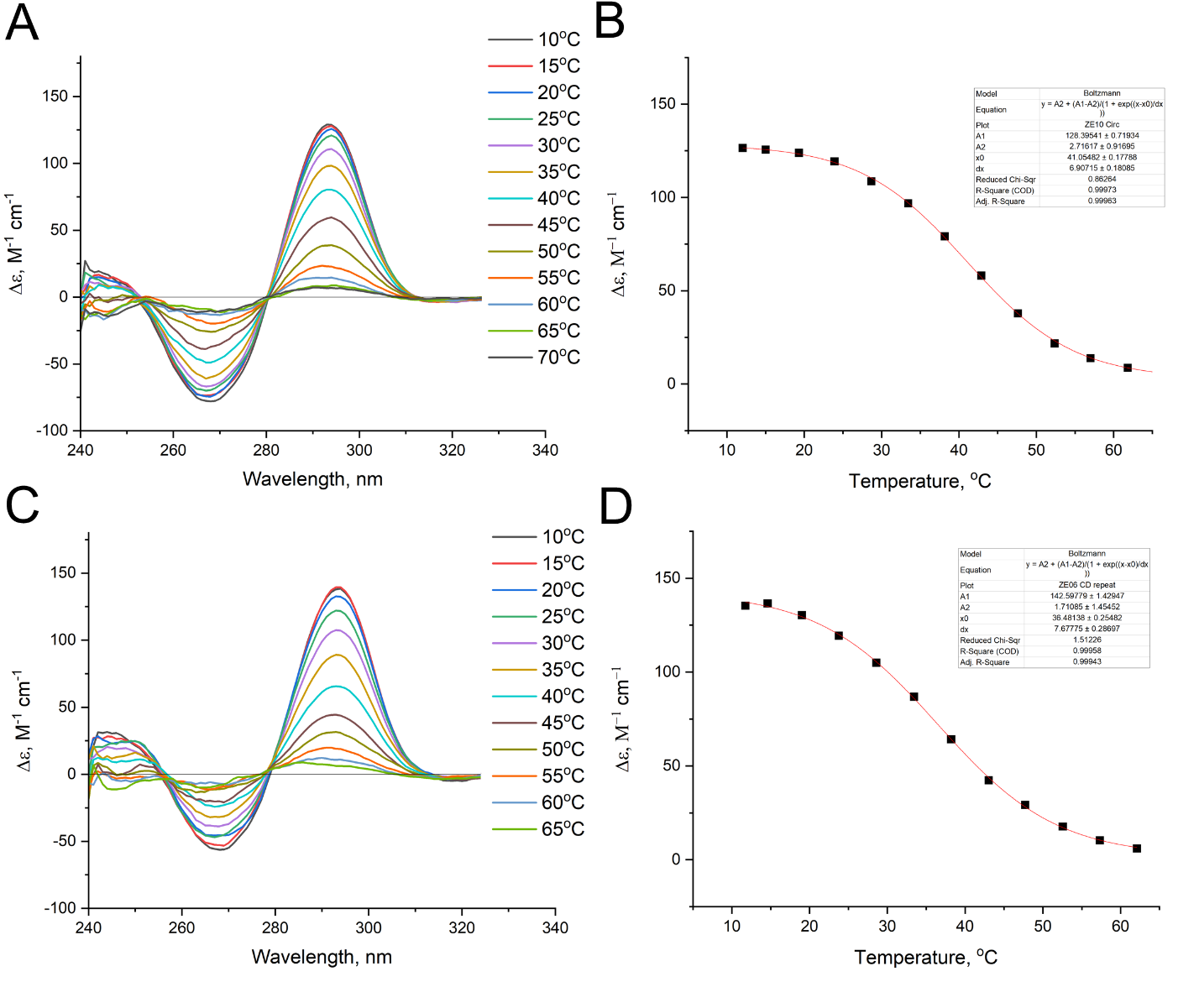

Figure S14. Thermal stability of G-quadruplexes of T4,T13-HD1 (A) and T12,T13-HD1 (C) aptamers studied using circular dichroism spectroscopy. The melting profiles of T4,T13-HD1 (B) and T12,T13-HD1 (D) aptamers at a wavelength of 295 nm are provided.

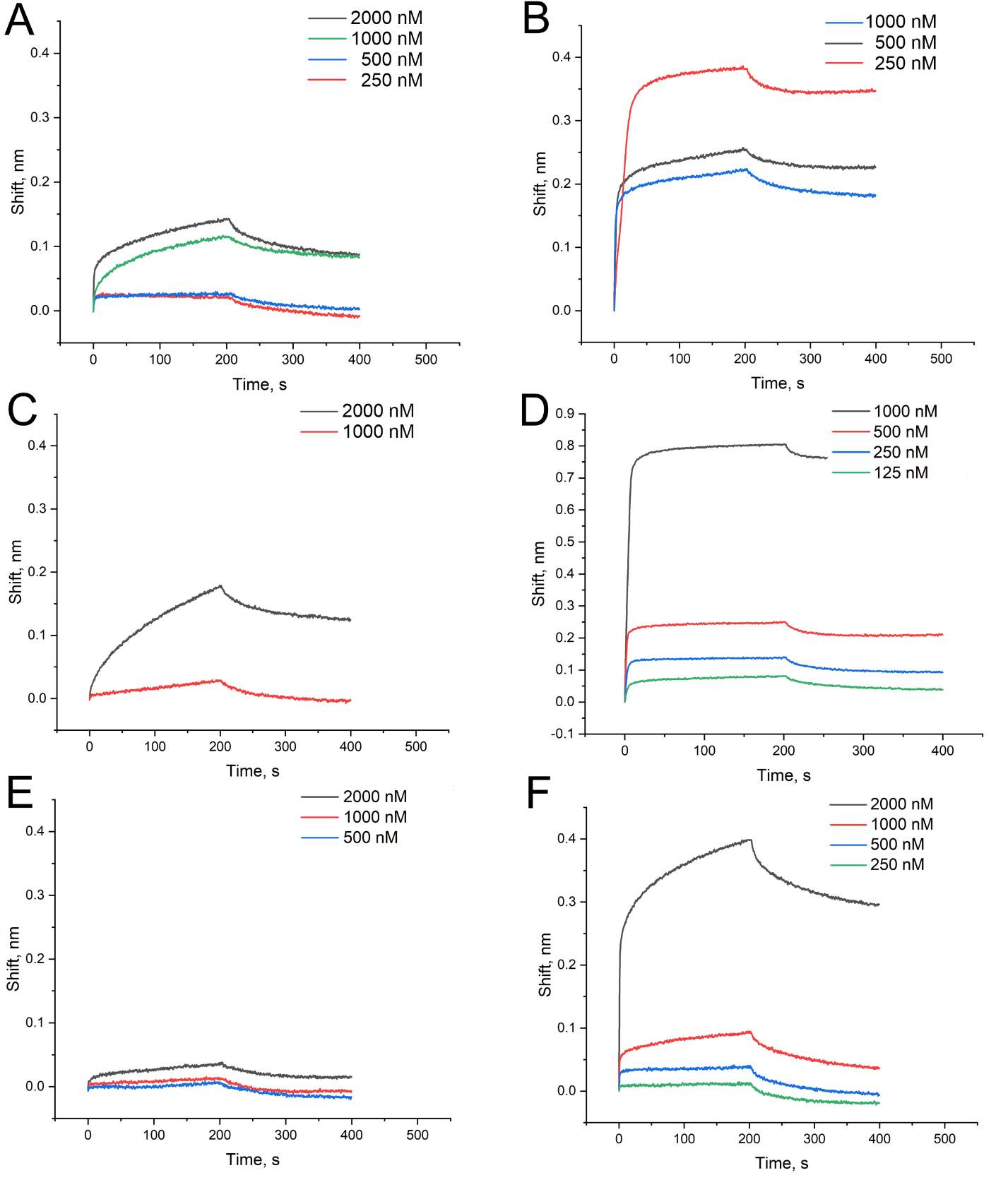

Figure S15. Biolayer interferometry sensorgrams for T3,T4-HD1 (A), T3,T12-HD1 (B), T3,T13-HD1 (C), T4,T12-HD1 (D), T4,T13-HD1 (E), T12,T13-HD1 (F) aptamers binding to immobilized thrombin. The aptamer-thrombin association stage was performed from 0 to 200 s; the aptamer-thrombin dissociation stage was performed from 200 to 400 s.

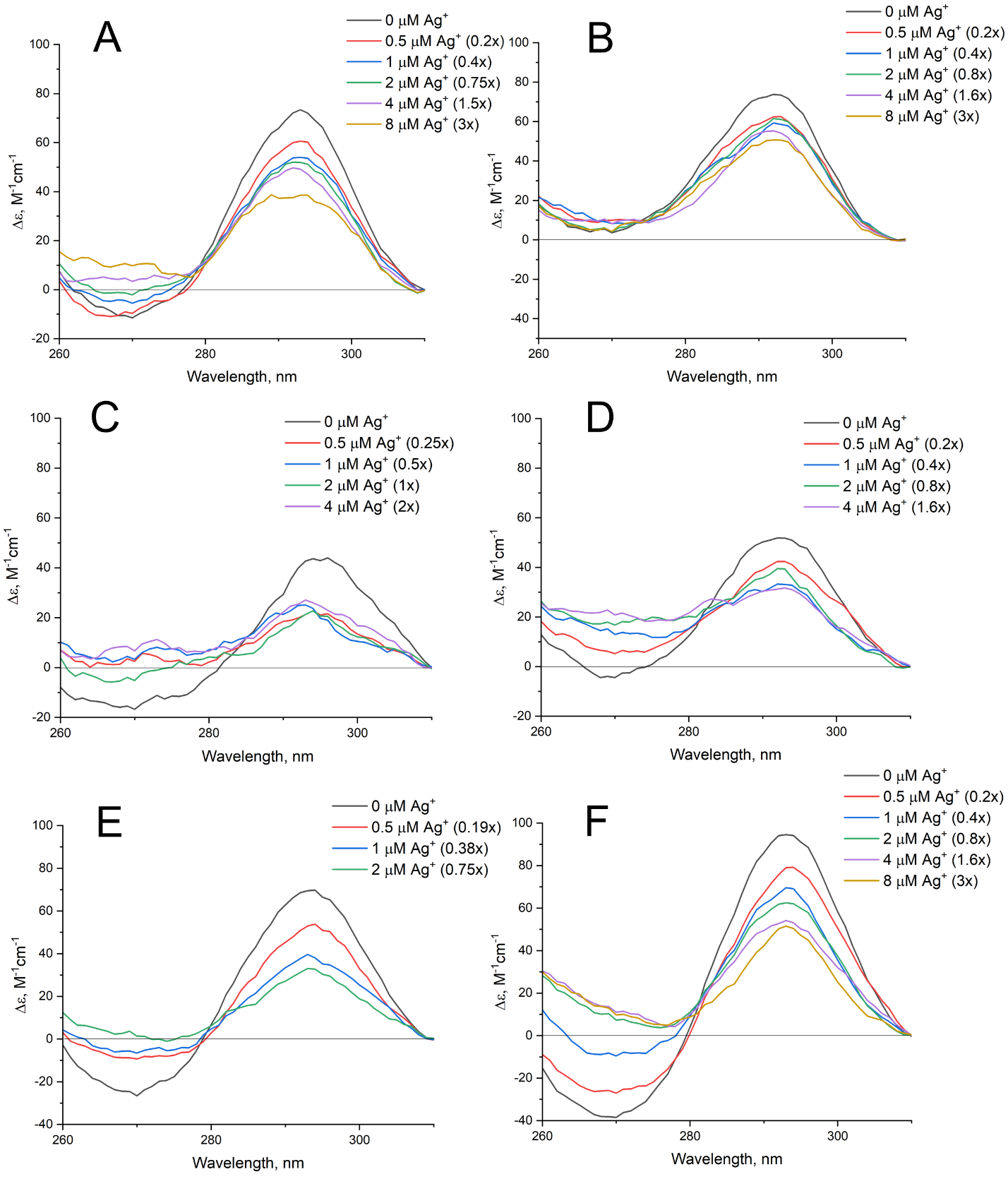

Figure S16. Circular dichroism spectra of aptamers with a dual oxo-εA modification in the presence of different amounts of Ag^+^ ions. T3,T4-HD1 (A), T12,T13-HD1 (B), and T3,T12-HD1 (C), T3,T13-HD1 (D), T4,T12-HD1 (E) and T4,T13-HD1 (F) aptamers were assembled in the buffer, mixed with Ag^+^ for 5 minutes and placed in the spectrometer.

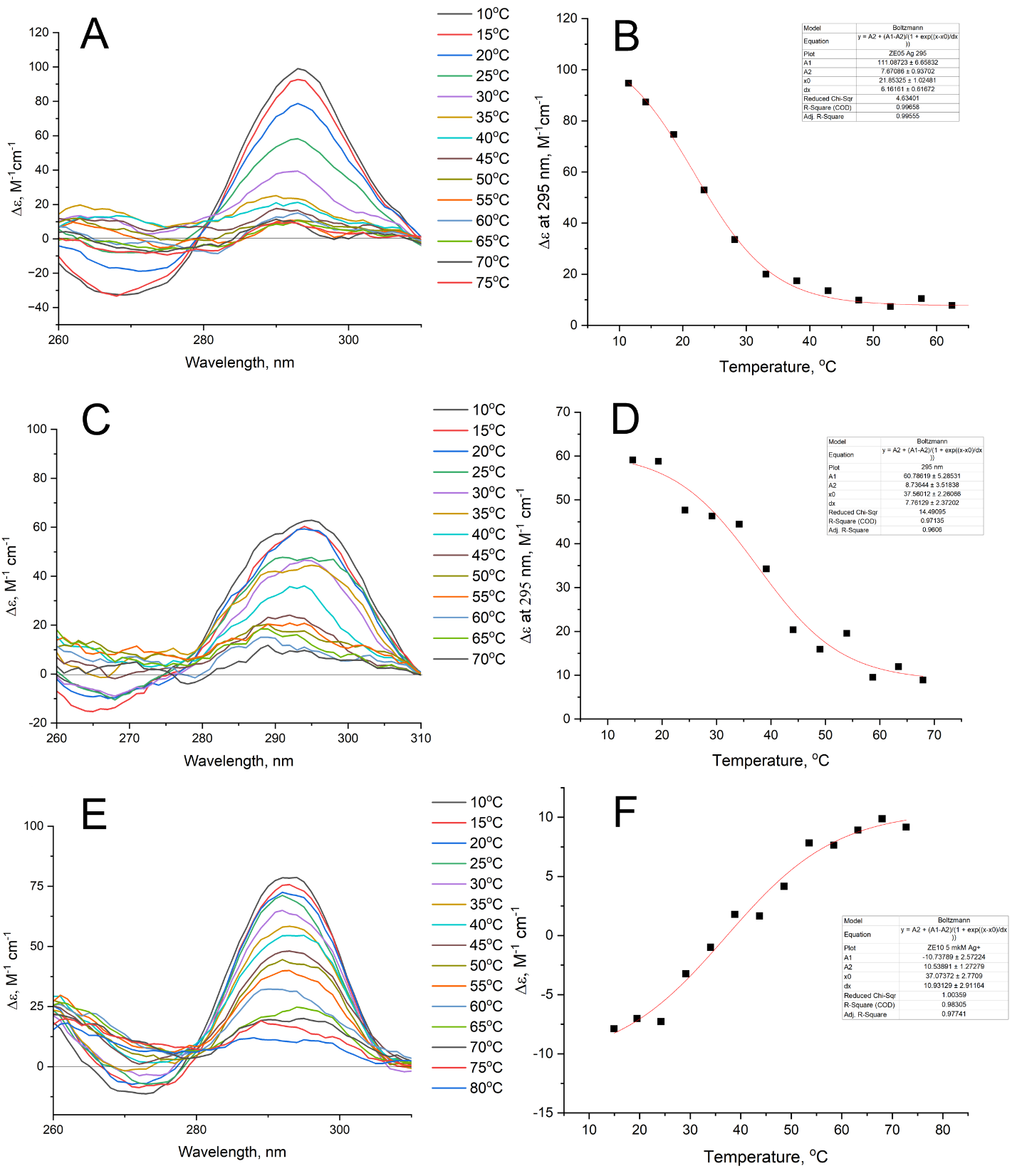

Figure S17. Temperature dependence of circular dichroism spectra of T3,T4-HD1 (A), T12,T13-HD1 (C), and T4,T13-HD1 (E) aptamers with 2-excess of Ag^+^ ions. The melting profiles at 295 nm are provided for T3,T4-HD1 (B) and T12,T13-HD1 (D) aptamers. The melting profile at 273 nm are provided for T4,T13-HD1 aptamer (F).

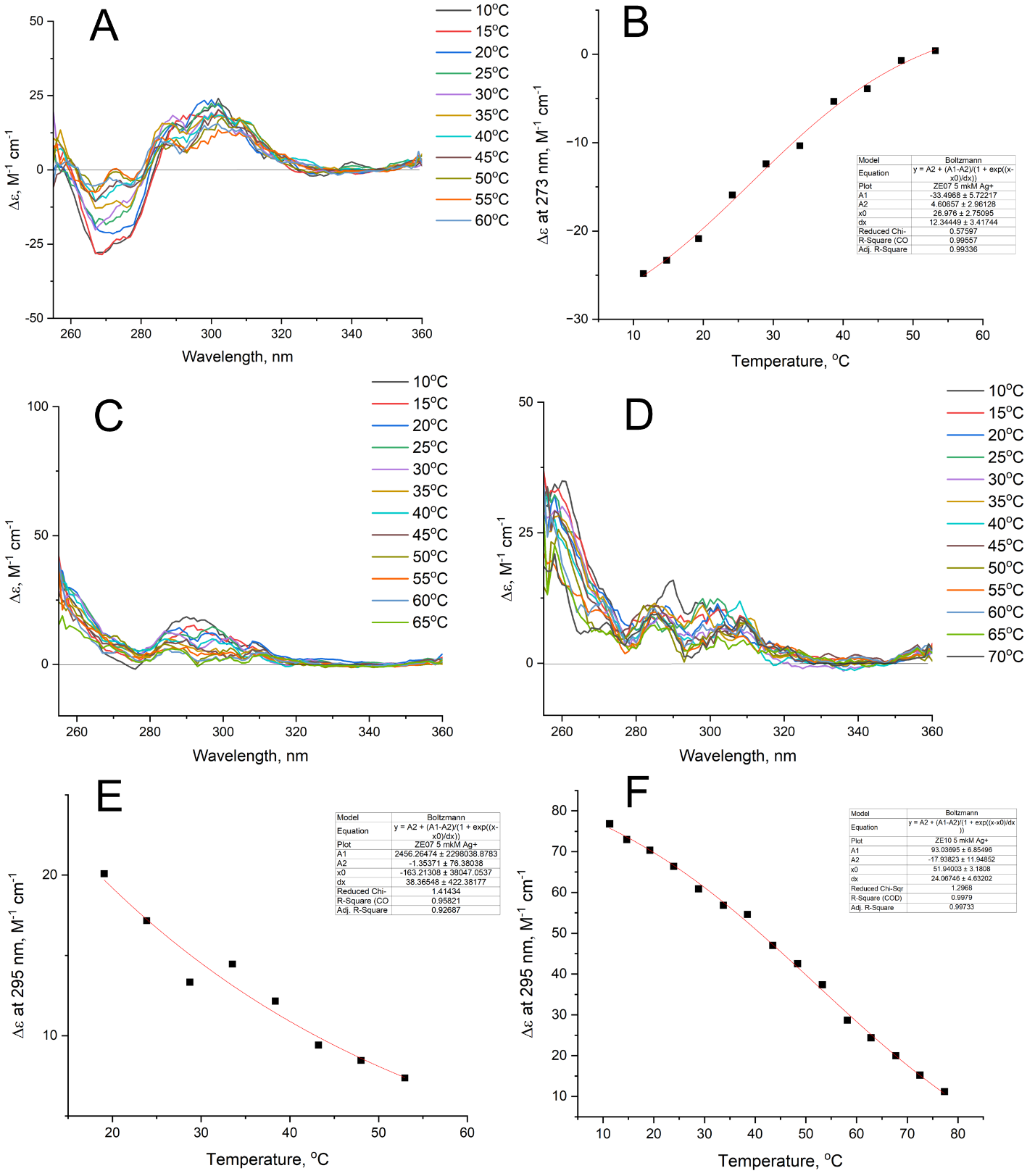

Figure S18. Temperature dependence of circular dichroism spectra of T3,T12-HD1 (A), T3,T13-HD1 (C), and T4,T12-HD1 (D) aptamers with 2-excess of Ag^+^ ions. The melting profiles at 273 nm are provided for T3,T12-HD1 aptamer (B). The melting profile at 295 nm are provided for T3,T12-HD1 (E) and T4,T13-HD1 (F) aptamers.

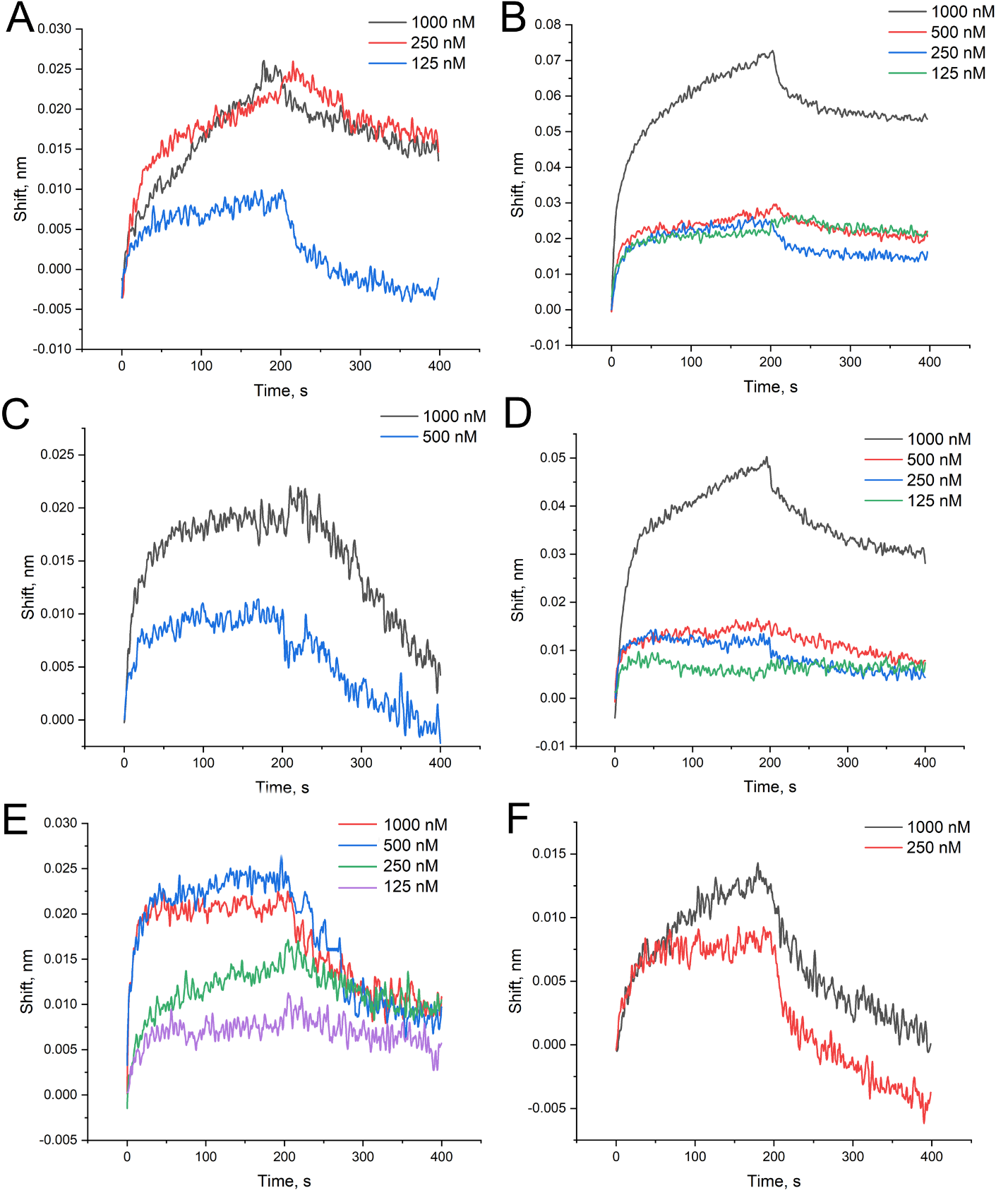

Figure S19. Biolayer interferometry sensorgrams for T3,T4-HD1 (A), T3,T12-HD1 (B), T3,T13-HD1 (C), T4,T12-HD1 (D), T4,T13-HD1 (E), T12,T13-HD1 (F) aptamers with 5-fold excess of Ag^+^ that bind immobilized thrombin. The aptamer-thrombin association stage was performed from 0 to 200 s; the aptamer-thrombin dissociation stage was performed from 200 to 400 s.
